# PI3K/mTOR is a therapeutically targetable genetic dependency in diffuse intrinsic pontine glioma

**DOI:** 10.1101/2023.04.17.537256

**Authors:** Ryan J. Duchatel, Evangeline R. Jackson, Sarah G. Parackal, Claire Sun, Paul Daniel, Abdul Mannan, Izac J. Findlay, Dilana Staudt, Zacary P. Germon, Sandra Laternser, Dylan Kiltschewskij, Padraic S. Kearney, M. Fairuz, B. Jamaluddin, Alicia M. Douglas, Tyrone Beitaki, Mika Persson, Elizabeth E. Manning, Heather C. Murray, Nicole M. Verrills, David A. Skerrett-Byrne, Brett Nixon, Susan Hua, Valdes-Mora Fatima, Maria Tsoli, David S. Ziegler, Murray J. Cairns, Eric Raabe, Nicholas A. Vitanza, Carl Koschmann, Frank Alvaro, Christopher V. Dayas, Christopher L. Tinkle, David D. Eisenstat, Ron Firestein, Sabine Mueller, Javad Nazarian, Jason E. Cain, Matthew D. Dun

## Abstract

Diffuse midline glioma (DMG), including tumors diagnosed in the brainstem (diffuse intrinsic pontine glioma – DIPG), are uniformly fatal brain tumors that lack effective pharmacological treatment. Analysis of pooled CRISPR-Cas9 loss-of-function gene deletion screen datasets, identified *PIK3CA* and *MTOR* as targetable molecular dependencies across DIPG patient derived models, highlighting the therapeutic potential of the blood-brain barrier penetrant PI3K/Akt/mTOR inhibitor paxalisib. At the human equivalent maximum tolerated dose, mice treated with paxalisib experienced systemic feedback resulting in increased blood glucose and insulin levels, commensurate with DIPG patients in Phase 1b clinical trials who experienced hyperglycemia/hyperinsulinemia. To exploit genetic dependences, but maintain compliance and benefit, we optimized a paxalisib treatment regimen that employed reduced dosing more frequently, in combination with the anti-hyperglycemic drug, metformin. Combining optimized dosing with metformin restored glucose homeostasis and decreased phosphorylation of the insulin receptor *in vivo*, a common mechanism of PI3K-inhibitor resistance, extending the survival of DIPG xenograft models. RNA sequencing and phosphoproteomic profiling of DIPG models treated with paxalisib identified increased calcium-activated PKC signaling. Using the brain penetrant PKC inhibitor, enzastaurin in combination with paxalisib, we synergistically extended the survival of orthotopic xenograft models, benefits further promoted by metformin; thus, identifying a clinically relevant DIPG combinatorial approach.

**Brief Summary:** Diffuse intrinsic pontine glioma is a lethal childhood brain tumor. Here we identify *PIK3CA* as a genetic dependency targeted by the brain penetrant pan-PI3K-inhibitor paxalisib.

## Introduction

Pediatric and adolescent diffuse midline glioma (DMG), including diffuse intrinsic pontine glioma (DIPG), are universally fatal high-grade gliomas (HGGs) diagnosed in the midline structures of the brain. DIPG is responsible for more brain tumor-related deaths in children than any other cancer (1). Palliative radiotherapy is the only approved treatment, with median overall survival of 9-11 months post-diagnosis (2, 3).

Global hypomethylation of histone H3 at lysine 27 (H3K27me3) is the biological hallmark feature of DIPG, leading to loss of gene silencing, promoting pro-oncogenic transcriptional programs for which there are no approved treatments (4, 5). Global loss of H3K27me3 is driven by recurring mutations in histone H3 genes including, *HIST1H3B/C* or *H3F3A* (4, 6), or through overexpression of the EZH inhibitory protein (EZHIP) (7), both of which inhibit the catalysis of H3K27 trimethylation by the polycomb repressive complex 2 (PRC2). The recent World Health Organization’s 5^th^ Classification of Central Nervous System Tumors designates DMG as ‘H3-K27-altered’ indicating that global hypomethylation of H3K27 is seen in the majority of patients with DMG (8). Here, we will use the term DIPG to collectively refer to both H3 wildtype (WT) and H3 K27M mutant diffuse pontine gliomas. H3-alterations in DIPG are instigating mutations, but are accompanied by an obligate partner mutation in a signaling gene (*PDGFRA, ACVR1, PIK3CA, PIK3R1, EGFR*), and/or a tumor suppressor gene (*TP53, PPM1D, PTEN, BCOR*), with co-occurrence of either necessary to induce gliomagenesis *in vivo* (9, 10). Co-segregation of discrete components of the PI3K/Akt/mTOR signaling axis are recognized as recurrent molecular drivers of H3-altered gliomas (3), with recurring mutations or amplifications in *PDGFRA* driving constitutive activity of the PI3K signaling axis (11). Activated PI3K/Akt/mTOR signaling drives angiogenesis, cancer cell metabolism, growth, and survival (10), highlighting the potential of therapies that show activity in the central nervous system (CNS) and target this oncogenic signaling axis for the treatment of DIPG.

Targeting PI3K/Akt/mTOR is a treatment paradigm that has been vigorously tested across almost all cancer types (12). Although there are >40 different inhibitors in various stages of clinical development, only mTOR inhibitors including temsirolimus (13) and everolimus (14), and PI3K inhibitors idelalisib and copanlisib (15), have gained FDA approval as anti-cancer therapies; however, these all show limited activity in the CNS.

The CNS penetrant, pan PI3K/Akt/mTOR (p110α, p110β, p110δ and p110γ) inhibitor paxalisib (formerly GDC-0084), was developed for the treatment of glioblastoma, as ∼80% of cases harbor recurring mutations and amplification in genes mapping to the PI3K signaling axis (16). Specifically optimized to cross the blood-brain barrier (BBB) (17), paxalisib has completed dose escalation and maximum tolerated dose (MTD) in DIPG clinical trials, identifying a MTD of 27 mg/m^2^/day (NCT03696355) (18). These studies followed human trials in adults with recurrent high-grade gliomas (HGGs), where paxalisib showed brain penetration at clinically relevant concentrations, with 40% of patients achieving stable disease (19). However, treatment-induced transient hyperinsulinemia is a major driver of reduced efficacy of PI3K/Akt/mTOR inhibitors, promoting glycogen breakdown in the liver and inhibition of glucose uptake in skeletal muscle and adipose tissues, resulting in hyperglycemia (20, 21). Hence, patients experience compensatory insulin release from the pancreas to restore normal glucose homeostasis, promoting side effects and insulin feedback pathways that reactivate PI3K/AKT/mTOR signaling in tumors, particularly when continuous PI3K inhibition is attempted in isolation (22).

The complex and heterogeneous somatic, epigenetic, and clonal landscapes of DIPG render monotherapeutic approaches unlikely to promote long-term survival (23, 24). Therefore, combination strategies that synergize and exploit the unique biological features of DIPG are desperately needed. Here, we have analyzed a targeted CRISPR-Cas9 gene deletion dataset and identified the dependency of DIPGs on PI3K/Akt/mTOR for the transmission of oncogenic signals. Furthermore, we have addressed the therapeutic limitations of paxalisib-induced transient hyperinsulinemia using dose optimization alone and in combination with metformin (18). Using a multiomic sequencing strategy including transcriptomics and quantitative phosphoproteomics of DIPG cells, we have identified increased calcium induced protein kinase C (PKC) signaling following paxalisib exposure, suggesting a combined therapeutic vulnerability. In this study we address the intrinsic neoplastic sequela of DIPG/DMG, by combined targeting of PI3K/Akt/mTOR using paxalisib, compensatory PKC signaling using enzastaurin, coupled with strategies to manage treatment-related side-effects and reduced efficacy using metformin. This is a clinically relevant and feasible combination strategy for the treatment of DMG patients to be studied in clinical trials.

## Material and Methods

### DIPG genetic dependency data

The genetic dependency data were derived from the Victorian Paediatric Cancer Consortium’s Childhood Cancer Model Atlas (CCMA) Data Portal that contains dependency data on 352 genes across 38 DMG cell line models (25).

### Cas9 Cell Generation and sgRNA design for PIK3CA knockdown

A total of 2 x 10^5^ cells were seeded in a 6-well plate and incubated overnight. Cells were then replenished with fresh complete media containing 5 µg/mL polybrene (ThermoFisher Scientific). A 250 µL aliquot of lentiviral cocktail containing either Lenti-Cas9-Blast plasmid (SU-DIPG-XIII; Addgene, Watertown, MA, USA) or Lenti-Cas9-2A-Blast (SU-DIPG-XXXVI; Addgene) was supplemented into the cell media and incubated for 72 h. Transduced cells were selectively maintained in complete media containing 10 µg/mL blasticidin (Jomar Life Research, Scoresby, Victoria, Australia) for at least 7 days. *PIK3CA* and non-targeting control (NTC) single guide RNA (sgRNA), cloned into the U6-gRNA/hPGK-puro-2A-BFP vector, were obtained from the Human Sanger Whole Genome Lentiviral CRISPR Library (ThermoFisher Scientific). The gRNA sequence for *PIK3CA* was 5’-GCAAATAATAGTGGTGATCTGG-3’, and the non-targeting control (NTC) 5’-CCCAACTTCAACACCAATCT-3’. A total of 5 × 10^5^ Lenti-X HEK29T were seeded in 6-well plates and the following day were transfected with sgRNA plasmids along with the viral packaging plasmids, psPAX-D64V (Addgene) and pMD2.G (Addgene) using Lipofectamine^TM^ LTX Reagent with PLUS^TM^ reagent as per the manufacturer’s recommendations. Transfection media was replaced with fresh media after 6 h and incubated for a further 72 h prior to collection of virus-containing media. Viral media was added to 2 × 10^5^ Cas9-expressing DIPG cells in a 6-well plate in the presence of 1 µg/mL polybrene, centrifuged at 800 x g for 30 min and then incubated for 72 h. Selection of transduced cells using 2 µg/mL of puromycin in fresh media was performed until non-transduced control cells were dead. Heterogenous cell lines were maintained in 2 µg/mL puromycin. For the establishment of single cell clones from the heterogenous population, single BFP positive cells were sorted in 96-well plates containing a 1:1 mixture of conditioned media and fresh media. Single cell clones were expanded and screened using immunoblotting to identify clones with reduced or absent target protein.

### High-throughput drug screening

Cellular growth and proliferation of DIPG neurosphere cell lines was determined using a resazurin growth and proliferation assay, as previously described (26).

### Western Blotting

Protein was extracted from DIPG cells using RIPA buffer as per manufacture’s recommendations and previously described (27). BCA quantification was performed using a Pierce BCA Protein Assay Kit (ThermoFischer Scientific) according to the manufacturer’s instructions. Primary antibodies were incubated overnight at dilutions described in Supplementary Table 1. Secondary horseradish peroxidase (HRP) conjugated antibody (Bio-Rad, Hercules, CA, USA) was used at a dilution of 1:5000 (antibodies in Supplementary Table S1). Labeled protein bands were imaged using enhanced chemiluminescence (Merck) in combination with a Chemidoc MP Imaging System (Bio-Rad) and data were analyzed using ImageLab software.

### Next generation sequencing analysis

Genomic profiling was performed on 16 DMG neurospheres using an Illumina TruSight Oncology 500 (TSO500, Illumina, CA, USA) next-generation sequencing (NGS) assay as previously described (28). Following FASTQ validation, sequences were aligned to the hg19 genome (29). PISCES was used to perform somatic variant calling to identify variants at low frequency, which are filtered when error rates do not meet quality thresholds (30). GEMINI was used to perform local INDEL realignment, paired-read stitching, and read filtering to identify insertion and deletion events as well as further improve variant calling results (31). The CRAFT copy number (CN) variant caller performed amplification, reference, and deletion calling for target copy number variation (CNV) genes within the assay (28). The CRAFT software component counts the coverage of each target interval on the panel, performs normalization, calculates fold change values for each gene, a quality control Q-score and determines the CNV status for each CNV target gene. Illumina Annotation Engine Nirvana software performed annotation of small variants, calculating tumor mutational burden and microsatellite instability for each sample (32). Finally, the MANTA fusion caller was utilized to discover, assemble, and score large-scale gene fusions (33). TSO500 contamination detection was implemented to identify contaminated samples by examining a combination of contamination p-value (p-score) and scores determined by error rate per sample and total read depth of the nucleotide position. VCF files for each sample produced by TSO500 were analyzed using the Helium application, allocating a Helium Score (a measure of likely pathogenicity of the variant based on its appearance in somatic mutation databases: COSMIC, Clinvar, CGC and CADD), and Global Max Allele Frequency (frequency of that variant within the global population), to each variant (34). Variants with a Global Max Allele Frequency of <1e^-5^ were predicted to be somatic unless present in a matched germline sample. CNVs for each gene were grouped into categorical states (copy number deletion [CN<1.32], copy number loss [CN between 1.32 and 1.7], no change [CN between 1.7 and 3], copy number gain [CN between 3 and 7] and copy number amplification [CN>7]) based on categories from genomic studies (3, 35). Predicted somatic variants and CNVs with a Q-score >125 in genes belonging to the PI3K/AKT/mTOR signaling pathway were plotted in an oncoprint using the ComplexHeatmaps package (36).

### Bulk RNA barcode (BRB) sequencing

BRB-sequencing (henceforth referred to as RNA-seq) was performed as previously described (37). SU-DIPG-VI neurospheres were seeded in 6-well plates (500 000 cells per well) and treated with paxalisib at IC_50_ (Supplementary Table S2). Cells were collected at time points 0, 6 and 12 h post treatment. BRB-seq library was prepared with a Mercurius BRB-seq kit for 96 samples (Alithea Genomics, Manual v.0.1.61) (38). RNA library quality was assessed using Agilent TapeStation (fragments between 300–1000 bp needed) and quantified using a Qubit High Sensitivity Assay. Pooled samples were sequenced to a depth of 441.4 million reads using the NextSeq v2.5 High Output (75 cycles) on the Illumina NextSeq 500 (Illumina). Volcano plots of the differential expression analysis of each treatment relative to DMSO control were constructed, and the threshold of significance for each gene was set to a false-discovery rate adjusted p-value (FDR) of less than 1 and an absolute log_2_ fold change greater than 1 (39). For heatmap visualization and clustering, datasets of each treatment along with control were normalized using variance stabilizing transformation from DESeq2, and batch effects were removed using the remove batch effect function in limma (40).

### High-resolution quantitative phosphoproteomic profiling

SU-DIPG-XXXVI neurospheres were treated with paxalisib and subjected to proteomic and phosphoproteomics analysis as previously described (41–43). SU-DIPG-XXXVI neurospheres were treated with paxalisib at IC_50_ (Supplementary Table 2) for 6 h. Tryptic peptides from each sample were prepared as described (42, 44, 45) individually labeled using tandem mass tags (TMT) and phosphopeptides were isolated from the proteome using titanium dioxide (44) and immobilized metal affinity chromatography before offline hydrophilic interaction liquid chromatography (HILIC) (26, 42, 43). Liquid chromatography (LC) tandem mass spectrometry (MS/MS) was performed using a Q-Exactive Plus hybrid quadrupole-Orbitrap MS system, coupled to a Dionex Ultimate 3000RSLC nanoflow HPLC system as described (42, 43).

### Pharmacokinetic analysis

Paxalisib was diluted in 1% methylcellulose cp15/0.2% Tween 80 and administered via oral gavage to 6-8-week-old, female, NSG mice at either 5 mg/kg, 10 mg/kg or two 5 mg/kg doses 12 h apart (5 mg/kg/b.i.d.). After 1, 6 or 24 h post treatment, mice were sacrificed by CO_2_ euthanasia. Immediately following, blood was extracted via cardiac puncture, and brains collected. Blood plasma was separated via standard centrifugation techniques and frozen at – 80°C. Brainstem, thalamus, and prefrontal brain regions were dissected prior to snap freezing in liquid nitrogen. Brain tissues were homogenized using Lysing Matrix beads in a FastPrep-24™ 5G system (MP Biomedicals) at the factory recommended settings. Paxalisib was extracted from plasma and homogenized brain tissues using a protein precipitating mixture composed of 90% v/v acetonitrile, 10% v/v ethanol and 0.1% v/v glacial acetic acid. The supernatant was separated and collected following centrifugation. The supernatants were then analyzed using a Nexera X2 UHPLC system (Shimadzu) coupled to a QTRAP 6500 System (SCIEX) via multiple reaction monitoring (MRM) using 282.0 and 240.1 transitions of the 367.2 m/z precursor mass as described (46). Quantitative analysis was conducted using MultiQuant Software (SCIEX) against a calibration curve for paxalisib over the concentration range of 0.061 – 1000.0 pg/μL (46).

### Tissue pharmacodynamics

A single cell suspension of SU-DIPG-XIII-Pons* neurospheres was injected into the pontine region of 6-8 week, female, NSG mice under isoflurane anesthesia. Stereotactic coordinates were 0.8 mm to the right of midline, 0.5 mm posterior to lambda and 5 mm deep using 500,000 cells in 2 µL. Two weeks post xenograft, mice were treated with as single dose (paxalisib) of 5 mg/kg, 10 mg/kg or two 5 mg/kg doses 12 h apart (5 mg/kg/b.i.d.), by oral gavage, before being sacrificed by CO_2_ euthanasia, 6, 12, or 24 h later. Immediately following euthanasia, mice were transcardially perfused with saline, brains removed, brainstem and prefrontal cortex dissected and snap frozen in liquid nitrogen. Tissue was homogenized on ice using a Dounce homogenizer in RIPA protein extraction buffer containing protease and phosphatase inhibitors. Homogenized samples were sonicated 5 × 20 sec at 4°C, before being clarified through centrifugation at 25,000 × *g* for 30 min at 4°C. Samples were then subjected to standard Western blotting techniques as previously described (27) using primary antibodies (Supplementary Table S1).

### Blood glucose and C-peptide analysis

Mice were sacrificed 4 h following final treatment and blood was collected via cardiac puncture for measurement of blood glucose and C-peptide. Blood was applied to glucose test strips (Abbott Freestyle) and levels of blood glucose were measured using an Abbott FreeStyle Optium Neo (Abbott Freestyle). Blood was centrifuged (2,500 × *g*, 15 min, 4°C) and plasma supernatant was stored at –80°C. A Mouse C-peptide ELISA (ALPCO), was then used to quantify C-peptide levels in the plasma, as per manufacturer’s recommendations.

### Orthotopic xenograft survival

A single cell suspension of SU-DIPG-XIII-P*, HSJD-DIPG-007, RA-055, or UON-VIBE5 neurospheres were injected into the pontine region of 6-8 week old, female, NSG mice under isoflurane anesthesia. Stereotactic coordinates were 0.8 mm to the right of midline, 0.5 mm posterior to lambda and 5 mm deep using 500,000 cells (SU-DIPG-XIII-P*), 300,000 cells (HSJD-DIPG-007), 200,000 cells (RA-055) or 300,000 cells (UON-VIBE5) in 2 µL HBSS. For all animal studies, animals were randomly allocated to treatment group. SU-DIPG-XIII-P* xenografted mice commenced treatment 7 days post xenograft, and HSJD-DIPG-007 and RAO55 xenograft mice commenced treatment 21 days post xenograft. UON-VIBE5 xenografted mice commenced treatment 35 days post xenograft after initial observation of DIPG symptoms in a subset of tumor bearing mice. To assess the monotherapeutic efficacy of paxalisib, mice were treated with 5 mg/kg daily, 10 mg/kg daily or 5 mg/kg/b.i.d., paxalisib by gavage in 1% methylcellulose cp15/0.2% Tween 80, every 5 days, continuously. To assess the combination effects of paxalisib and targeted therapies, 5 mg/kg twice daily dosing of paxalisib was combined with metformin (175 mg/kg, day, gavage), vandetanib (25 mg/kg, daily, gavage), ribociclib (75 mg/kg, daily, gavage) or enzastaurin (100 mg/kg, daily, gavage), in 1% methylcellulose cp15/0.2% Tween 80 continuously or for a maximum of five weeks. Mice were euthanized at an ethical end point, including when exhibiting signs of neurologic decline, such as ataxia, circling or head tilting, with or without 20% weight loss. Overall survival was determined via the Kaplan-Meier survival analysis. GraphPad Prism Version 9.1.0. was used for in vivo statistical analyses using the Mantel-Cox test. In all cases, values of *p<0.05* were regarded as being statistically significant compared to vehicle, and *p<0.01* considered the threshold for combination synergism compared to monotherapies as described (47).

### Immunohistochemistry

Immunohistochemical staining was performed by the Hunter Medical Research Institute (HMRI) Core Histology Facility. Immunohistochemistry optimization and staining services were provided by The University of Newcastle’s ‘NSW Regional Biospecimen Services’. Staining was performed on the Discovery Ultra Benchmark Immunohistochemistry Automated Platform (Roche). Briefly, antigen retrieval was conducted at 95°C for 32 mins, blocked at 37°C for 12 mins before addition of primary antibodies per Supplementary Table 1. Anti-Rabbit HQ was applied for 16 mins at 37°C followed by tertiary of HRP HQ for 16 mins at 37°C and developed using diaminobenzidine (DAB). Slides were counterstained using freshly filtered haematoxylin for 10 seconds, before being washed and dehydrated per standard procedures (Supplementary Table S1).

### Data availability statement

The phosphoproteomics data have been deposited to the ProteomeXchange via the PRIDE database. Data is available via ProteomeXchange with identifier PXD036114 (reviewer access username: password; oXb9NPYM). Raw and processed BRB-Seq files are available for download from the Gene Expression Omnibus (GEO) with accession number GSE211565 (reviewer access token: cfkpisckvzaxnkn).

### Statistical analysis

GraphPad Prism software (Version 9.1.0; La Jolla, CA, USA) was used to produce graphs and for statistical analysis of data. Unless otherwise stated, two sample, unpaired t-tests or one-way ANOVA were used to determine significant differences between groups. Event free survival analysis was performed using the Log-rank test. No data were excluded. Significant differences were detected in preliminary studies in our assays, prompting the use of minimum sample sizes for all *in vivo* experiments. *In vitro* experiments were performed at least 3 times each, per standard practices. Blinding was not performed in this study. Values shown are the mean ± SEM. Significance values: * = *p<0.05*, ** = *p<0.01*, *** = *p<0.001*, **** = *p<0.0001* are used throughout.

### Study approval

The use of human DIPG/DMG patient derived neurosphere cell lines was approved by the Human Ethics Research Committee, University of Newcastle (H-2018-0241), Human Ethics Research Committee, Monash University (HREC/17/MonH/323) and the Kantonale Ethikkommission Zurich (BASEC-Nr.2019-00615). All *in vivo* studies were approved by the University of Newcastle Animal Care and Ethics Committee (#A-2019-900 and #A-2020-004).

## Results

### Integrated CRISPR-Cas9 loss-of-function and drug screening predicts *PIK3CA* and ***MTOR* to be genetic dependencies in DMG**

To determine the importance of the expression of PI3K/Akt/mTOR genes in the transmission of oncogenic signals that promote the growth and proliferation of DIPG, we analyzed a CRISPR-Cas9 loss-of-function screen dataset performed on 38 DMG cell lines, representing all DMG H3-altered subtypes (25). Of the 13 genes mapping to the PI3K/Akt/mTOR signaling axis, strong genetic dependency is shown for *PIK3CA* and *MTOR* (Figure 1A). This was confirmed using two DIPG patient derived models that showed significantly diminished proliferation *in vitro* following the knockout/knockdown of *PIK3CA* (Supplementary Figure S1A). Conversely, DMG are not dependent on the negative regulator of PI3K-signaling, *PTEN*, and the positive Z-score is suggestive that inactivation of PTEN confers a growth advantage (Figure 1A).

**Figure 1:**
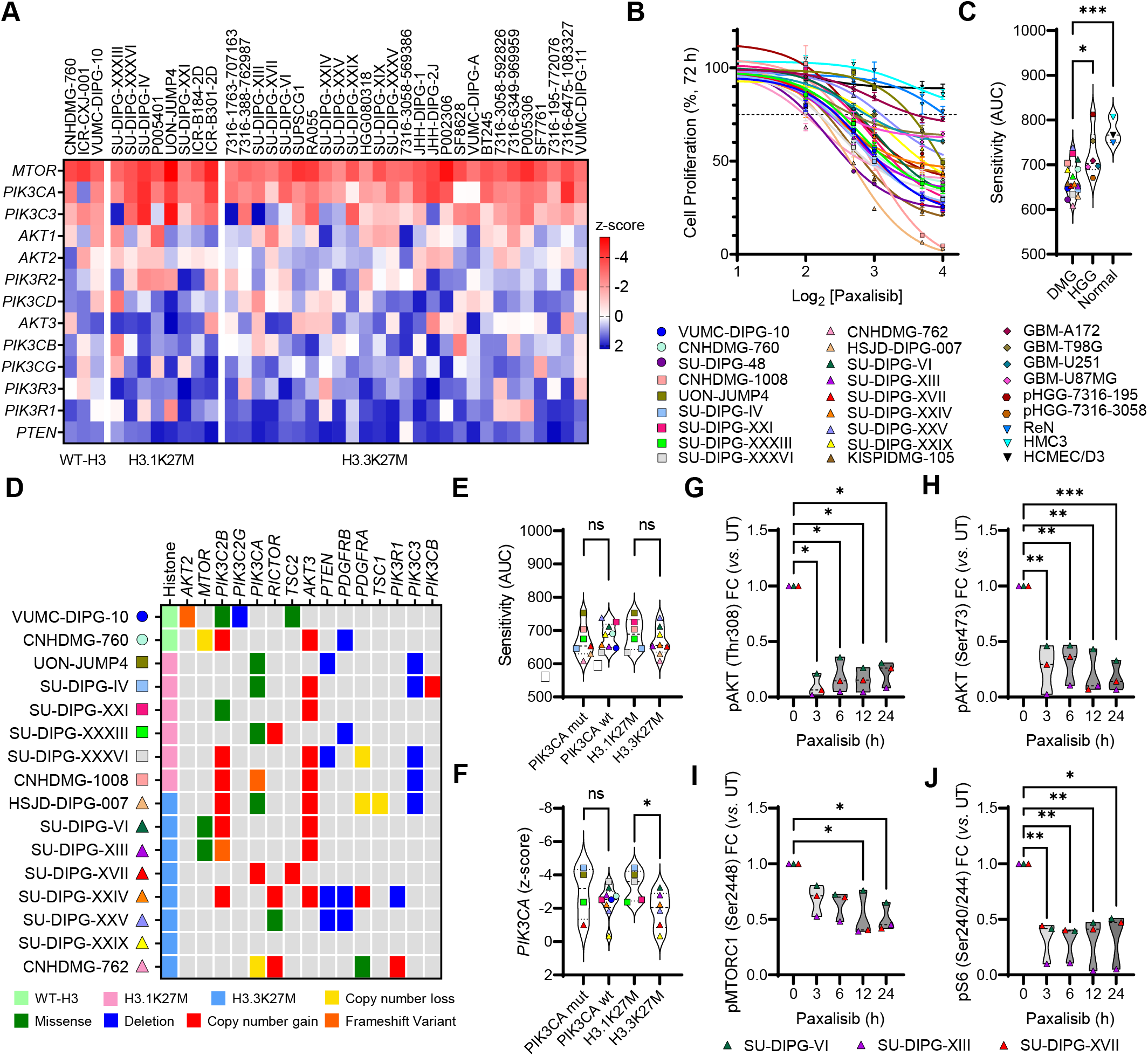
DMG patient derived cell lines are sensitive to paxalisib *in vitro*. (A) CRISPR-Cas9 loss-of-function screening across H3-altered G; wildtype-H3 (EZHIP) (n=3), H3.1K27M (n=8), H3.3K27M (n=27). (B) Sensitivity of DMG wildtype-H3 (EZHIP) (circles), H3.1K27M 3K27M (triangles), (n=18), GBM (diamonds) (n=4) and HGG (hexagons) (n=2) patient derived cell lines to paxalisib treatment as esazurin cell growth and proliferation assay. (C) Comparison of DIPG to HGG/GBM cell lines and normal cell lines (HCMEC/D3 rier (BBB) endothelial cells, HMC3 microglial cells and ReN neural progenitor cells) as measured by the sensitivity (area under the to paxalisib after 72 h (DIPG cell lines vs. HGG/GBM, *p*=0.0023 and normal *p=*0.0008, one-way ANOVA). (D) Oncoprint of PI3K DIPG cell lines (n=16) determined by TSO500 next generation sequencing. 12.5% harbored wildtype-H3 (EZHIP), 37.5% H3.1K27M K27M. (E) Area under the curve (AUC) comparison of paxalisib sensitivity between mutant versus wildtype *PIK3CA* DIPG cell lines mutation subgroups (*PIK3CA-*mut vs. *PIK3CA*-WT; H3.1K27M vs. H3.3K27M; one-way ANOVA, ns=not significant). (F) Analysis of tivity vs. *PIK3CA* z-score in *PIK3CA* mutant versus wildtype DIPG cell lines and H3K27M mutation subgroups (*PIK3CA-*mut vs. 3.1K27M vs. H3.3K27M; \**p*<0.05, one-way ANOVA, ns=not significant). (G-J) Quantitation of changes in phosphorylation of key R signaling proteins after treatment with 1 µM paxalisib for 24 h, in SU-DIPG-VI, SU-DIPG-XIII and SU-DIPG-XVII cells (n=3, one-*p<0.05*, \*\**p<0.01*, \*\*\**p<0.001*).

The high frequency, and hence evolutionary selection of recurring mutations in *PIK3CA*, *PIK3R1* and *PTEN* seen in DMG/DIPG (3, 10, 23) provided the impetus to target PI3K-signaling, upstream in the signaling cascade, rather than downstream targeting of mTOR directly. Analyzing the Cancer Dependency Map (DepMap) (48) additionally classified *PIK3CA* to be a strongly selective dependency in other cancer types, however *MTOR*, was identified to be a common dependency in both healthy and cancerous cells providing further justification for targeting PI3K in DIPG. Hence, we profiled proliferation of cells treated with the brain penetrant PI3K/Akt inhibitor paxalisib using DIPG, and non-midline pediatric HGG cell lines (glioblastoma) (Figure 1B). DIPG cells lines were significantly more sensitive to paxalisib than non-midline HGGs, with normal controls (HCMEC/D3 blood-brain barrier (BBB) endothelial cells, HMC3 microglial cells and ReN neural progenitor cells) resistant to treatment at high dose (Figure 1B–C). To determine whether recurring somatic alterations influence sensitivity to paxalisib, representative DIPG cell lines were subjected to next generation sequencing (NGS) (Figure 1D). Cell line models harbored H3-alterations commensurate with that seen in the DMG patient population (60:30:10, H3.3K27M: H3.1K27M: H3-WT) (3). Thirty percent of the models tested also harbored mutations in *PIK3CA*, 23% carried amplifications or mutations in *mTOR* or *RICTOR*, with the loss of PTEN identified in 25% of models (Figure 1D). No significant difference was seen in the sensitivity to paxalisib between DIPG neurospheres harboring WT or *PIK3CA*-mutations, or between H3.1K27M and H3.3K27M subtypes (Figure 1E).

By combining CRISPR-Cas9 *PIK3CA* dependency data and NSG data, we set out to determine the importance of recurring mutations on the level of *PIK3CA* dependency (Figure 1A, D). These analyses identified no difference in the level of *PIK3CA* dependence comparing *PIK3CA-*mutant verses WT-*PIK3CA* DIPG models (Figure 1F). However, H3.3K27M models were significantly more dependent on *PIK3CA* expression than H3.1K27M models (Figure 1F).

As activation of the PI3K pathway induces phosphorylation of downstream effector proteins, such as Akt and mTOR, we assessed protein expression and phosphorylation to determine whether abundance of pAkt (Thr308/Ser473), pMTORC1 (Ser2448) and pS6 (Ser240/Ser244) correlated with paxalisib sensitivity (Supplementary Figure S1B). Phosphorylation and hence activation of PI3K proteins was seen across DIPG models; however, the level of phosphorylation did not correlate with their *in vitro* sensitivity to paxalisib (Supplementary Figure S1C). Treatment of DIPG cell line models with paxalisib potently inhibited PI3K/Akt/mTOR phosphorylation, sustained for up to 24 h post-treatment *in vitro* (Figure 1G–J, Supplementary Figure S1D).

### Optimized *in vivo* dosing improved the pharmacokinetic and pharmacodynamic properties of paxalisib in the CNS

Historically, PI3K inhibitors have shown limited benefit for patients with CNS tumors due to their limited capacity to penetrate the BBB. Therefore, increased dosing is required to achieve brain concentrations sufficient to effectively suppress PI3K-signaling in brain tumors. This in turn, promotes PI3K-inhibitor-related side effects (rash, mucositis, neutropenia, and hyperglycemia), reducing patient compliance (49). First-in-human Phase I studies determined a MTD of 45 mg/day paxalisib in the adult recurrent HGG setting, with patients experiencing classical PI3K/mTOR–inhibitor related toxicities (19). These studies showed that at this dose, paxalisib crossed the BBB and had on-target effects. Importantly, oral low dose also showed good brain pharmacokinetic properties, and effective tumor growth inhibition even used once daily, with a C-max reached at 2 h post oral treatment (19). Subsequently, preliminary Phase 1B paxalisib safety and dose escalation studies in children with DIPG, determined a MTD of 27 mg/m^2^/day, in line with the adult MTD (based on a 60 kg adult), with patients also experiencing classical PI3K-related toxicities (18). Therefore, to establish *in vivo* CNS pharmacokinetics of paxalisib, we treated tumor naive NSG mice orally using the approximate mouse equivalent of the human MTD (10 mg/kg/day) (50), and reduced half-MTD once (5 mg/kg/day) or twice daily (5 mg/kg/b.i.d.) (Figure 2A–D). No significant loss of weight was seen across any of the dosing regimens following two weeks of treatment (Supplementary Figure S2A). Pharmacokinetic analysis showed increased plasma concentrations across all time points using 10 mg/kg/day compared to vehicle and 5 mg/kg once or twice daily (Figure 2A). This trend was also seen for 10 mg/kg/day compared to 5 mg/kg/day and 5 mg/kg/b.i.d. in brain tissues at some timepoints, including the prefrontal cortex (at 1 h, Figure 2B), thalamus (at 6 h post dosing, Figure 2C) and brainstem (at 1h, Figure 2D). However, in the brainstem, a significant increase in accumulation of paxalisib was seen for mice treated with 5 mg/kg/b.i.d., after 24 h, compared with 5 mg/kg/day, and a non-significant increase compared to 10 mg/kg/day (Figure 2D), in line with the 20.6-hour half-life of the drug (19).

**Figure 2:**
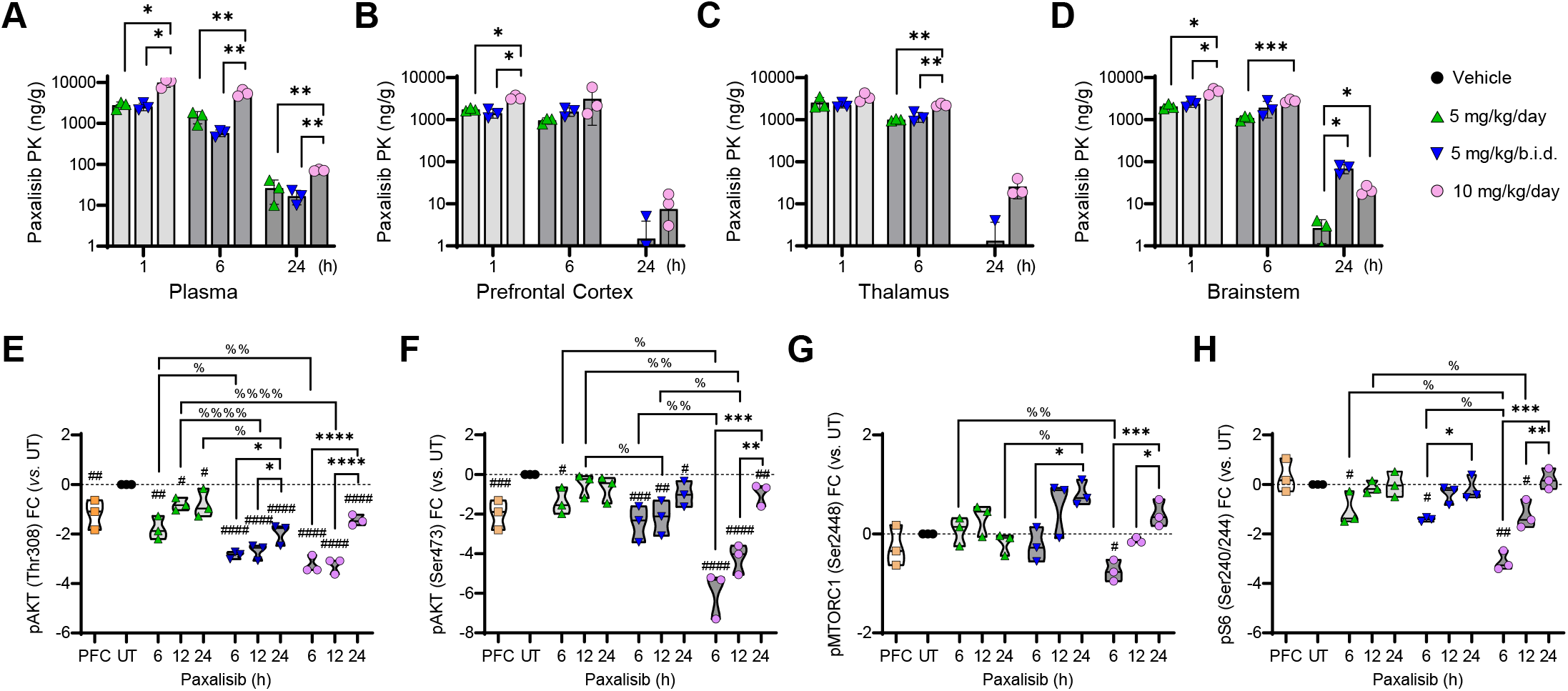
Paxalisib pharmacokinetics and pharmacodynamics *in vivo*. (A-D) Paxalisib pharmacokinetics and (E-H) pharmacodynamics fied dosing. (A-D) Concentration of paxalisib in (A) plasma, (B) prefrontal cortex (PFC), (C) thalamus and (D) brainstem, measured action monitoring mass spectroscopy (MRM) +/-treatment with paxalisib at 5 mg/kg/day, 5 mg/kg/b.i.d., or 10 mg/kg/day and r 1, 6 and 24 h (one-way ANOVA). (E-H) Phosphorylation analysis using SU-DIPG-XIII-P* tumor tissue treated with 5 mg/kg/day, 5 r 10 mg/kg/day paxalisib for two weeks and resected 6, 12 and 24 h post treatment (one-way ANOVA, *^/%^*p<0.05*, **^/%%^*p<0.01*, *1*, ****^/%%%%^*p<0.0001*, treated vs. untreated; ^#^*p<0.05*, ^##^*p<0.01*, ^###^*p<0.001*, ^####^*p<0.0001*).

To identify pharmacodynamic markers of successful PI3K/Akt/mTOR inhibition *in vivo* using these regimens, we engrafted SU-DIPG-XIII-P* patient derived DIPG cells into the pons of NSG mice. Tumors from SU-DIPG-XIII-P* xenograft mice were resected 28 days post-surgery after treatment with an acute dose of paxalisib at either 5 mg/kg/day, 5 mg/kg/b.i.d., or 10 mg/kg/day. Commensurate with *in vitro* analysis, the phosphorylation of pAkt (Thr308/Ser473) decreased in a dose dependent manner (Figure 2E–F, Supplementary Figure S2B). Treatment with 5 mg/kg/b.i.d., maintained suppression of PI3K signaling to a similar level to that of 10 mg/kg/day (Figure 2E–F, Supplementary Figure S2B). Rebound PI3K signaling was seen after 24 h, however, to a lesser extent using 5 mg/kg/b.i.d. (Figure 2E–F, Supplementary Figure S2B) and commensurate with the increased paxalisib accumulation seen in the brainstem of mice at 24 h using this regimen (Figure 2D). Although 5 mg/kg/b.i.d. significantly decreased phosphorylation of pAkt (Ser473) and phosphorylation of pS6 (Ser240/Ser244) compared to 5 mg/kg/day, 10 mg/kg/day was more effective than both reduced dosing regimens (Figure 2H, Supplementary Figure S2B).

### Paxalisib treatment modulates insulin and IP3 signaling in DIPG cell lines

To further our understanding of the anti-DIPG effects of paxalisib treatment, we performed RNA-seq using SU-DIPG-VI neurospheres following 6 and 12 h *in vitro* paxalisib exposure (at IC_50_ from Figure 1B, Supplementary Table S3-S5) and compared the results to DMSO vehicle treated controls. Analysis of both timepoints identified a total of 12,285 unique genes, quantitatively sequenced across samples (Supplementary Figure S3A), with 526 significantly downregulated transcripts (4.2% of total genes) and 454 significantly upregulated transcripts (3.7% of total genes), some of which were validated at the protein level via Western blotting (Supplementary Figure S3B). Ingenuity Pathway Analysis (IPA) was used to uncover the major canonical networks modulated by significant changes in transcript abundance following paxalisib treatment, and identified PI3K/Akt signaling, the insulin receptor, cholesterol biosynthesis, NRF2-mediated oxidative stress response, MYC, PPAR and EIF2 signaling (Figure 3A). Upstream regulator analysis identified increased PTEN, VEGF and CDK signaling (Figure 3B) and decreased regulation of insulin signaling (Figure 3C). Intriguingly, given the effect paxalisib has on systemic glucose homeostasis, the major transcription factor predicted to be upregulated by paxalisib treatment was *MLXIP*, the glucose-regulated factor of the Myc/Max/Mad superfamily (Figure 3D). Decreased upstream regulator analysis identified the SREBF1/2 transcription factors that control cholesterol homeostasis by regulating the transcription of sterol-regulated genes (Figure 3E). Altogether, potent modulation of insulin receptor signaling was seen in DIPG cells alongside that of increased activity of glucose-regulated pathways following paxalisib treatment *in vitro* (Figure 3A,C,E). It is important to note that insulin feedback is a well characterized mechanism of resistance to PI3K inhibitors *in vivo* (20), an important systemic consideration when testing paxalisib against DIPG *in vivo*.

**Figure 3:**
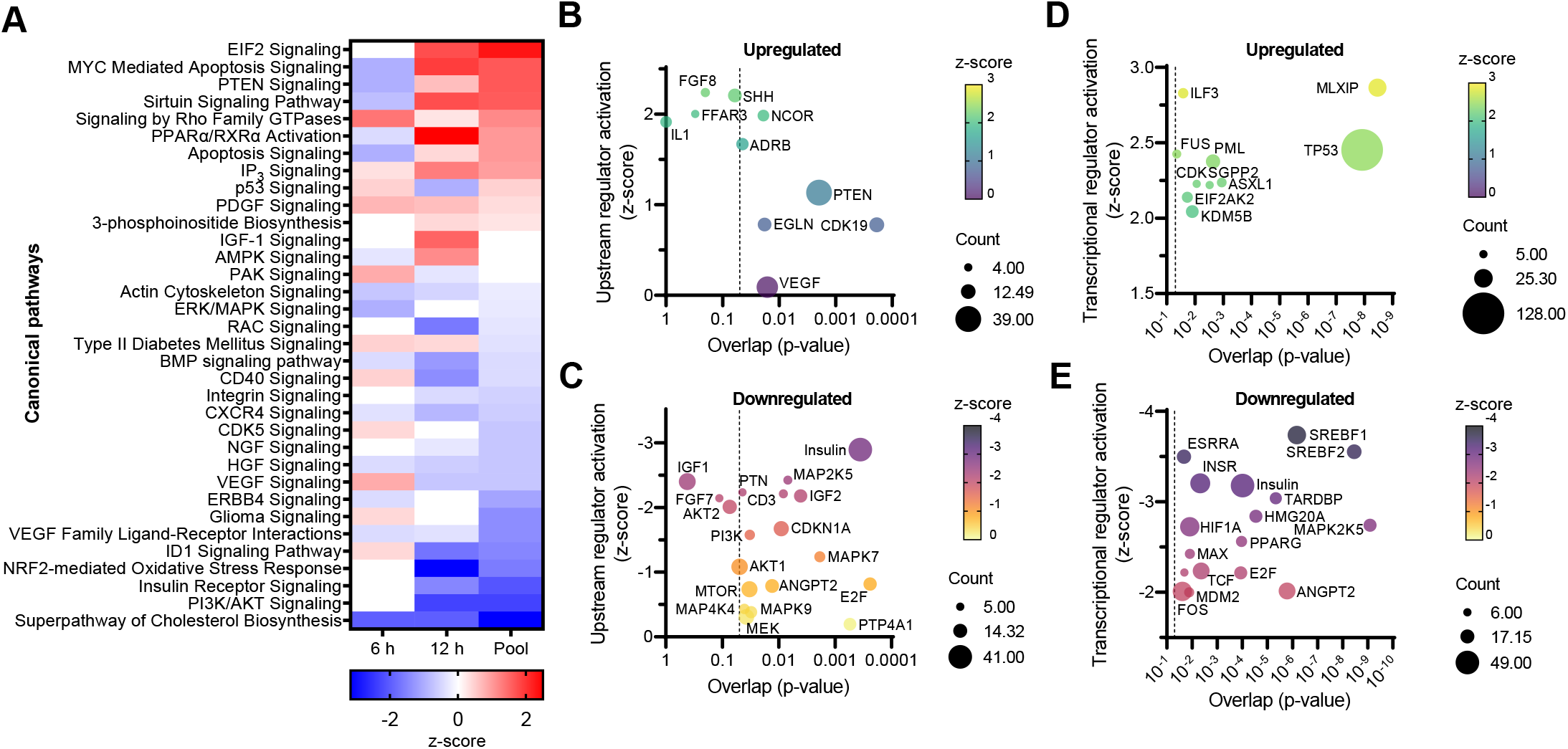
**Paxalisib treatment modulates insulin signaling in DIPG**. Bulk RNA barcoding and sequencing (BRB-seq) of SU-DIPG-VI following 6 µM paxalisib exposure. (A) Canonical pathway analysis of genes significantly altered by paxalisib exposure identified by Ingenuity sis (IPA). Activated pathways show a positive z-score in red, while inactivated pathways show a negative z-score in blue. Upstream sis of (B) significantly increased and (C) decreased expressed genes following paxalisib treatment, plotted using activation z-score, ze correlating to number of target molecules in dataset using IPA. Analysis of specific transcriptional regulators using significantly (D) wnregulated genes following of paxalisib treatment, plotted using activation z-score, *p*-value, and number of target molecules in IPA. one-way ANOVA, *^/%^*p<0.05*, **^/%%^*p<0.01*, ***^/%%%^*p<0.001*, ****^/%%%%^*p<0.0001*, treated vs. untreated; ^#^*p<0.05*, ^##^*p<0.01*, ^###^*p<0.001*, ^####^*p<0.0001*).

### Paxalisib treatment at the maximum tolerated dose (MTD) promoted hyperinsulinemia / hyperglycemia, rescued using half MTD alone and in combination with metformin

Grade 3 hyperglycemia was reported as the only dose limiting toxicity (DLT) for children treated with DIPG treated with paxalisib at 27 mg/m^2^ (18). Most frequently, grade 3 or 4 adverse events at MTD were rash (45%), neutropenia (36%), and hyperglycemia (20%), with the observed half-life of paxalisib determined at 20.6 ± 9 h in children with DIPG (18), similar to adult studies (19).

To address the reported hyperinsulinemia / hyperglycemia seen in clinical studies, SU-DIPG-XIII-P* pontine orthotopic xenograft mice were treated with modified paxalisib dosing regimens, alone and in combination with metformin (a commonly prescribed drug for type 2 diabetes, used to control blood glucose), daily for two weeks (5 on 2 off) and sacrificed 4 h post-last treatment to assess fasting blood glucose and C-peptide levels (surrogate measure of insulin levels). In line with previous results with DIPG patients, mice treated with the human-equivalent MTD of paxalisib (10 mg/kg/day) experienced significantly elevated blood glucose levels (∼200 mg/dL) (Figure 4A) and C-peptide levels 4 h post-treatment (Figure 4B). Mice treated with paxalisib at lower doses (5 mg/kg/day and 5 mg/kg/b.i.d.) still experienced increased blood glucose and C-peptide levels compared to vehicle controls; however, this occurred significantly less than those treated with 10 mg/kg/day (Figure 4A–B). Indeed, treatment with metformin significantly decreased blood glucose and C-peptide levels in mice treated with 10 mg/kg/day paxalisib, but not to the baseline levels, whereas metformin decreased both blood glucose and C-peptides levels back to vehicle controls in both 5 mg/kg regimens.

**Figure 4:**
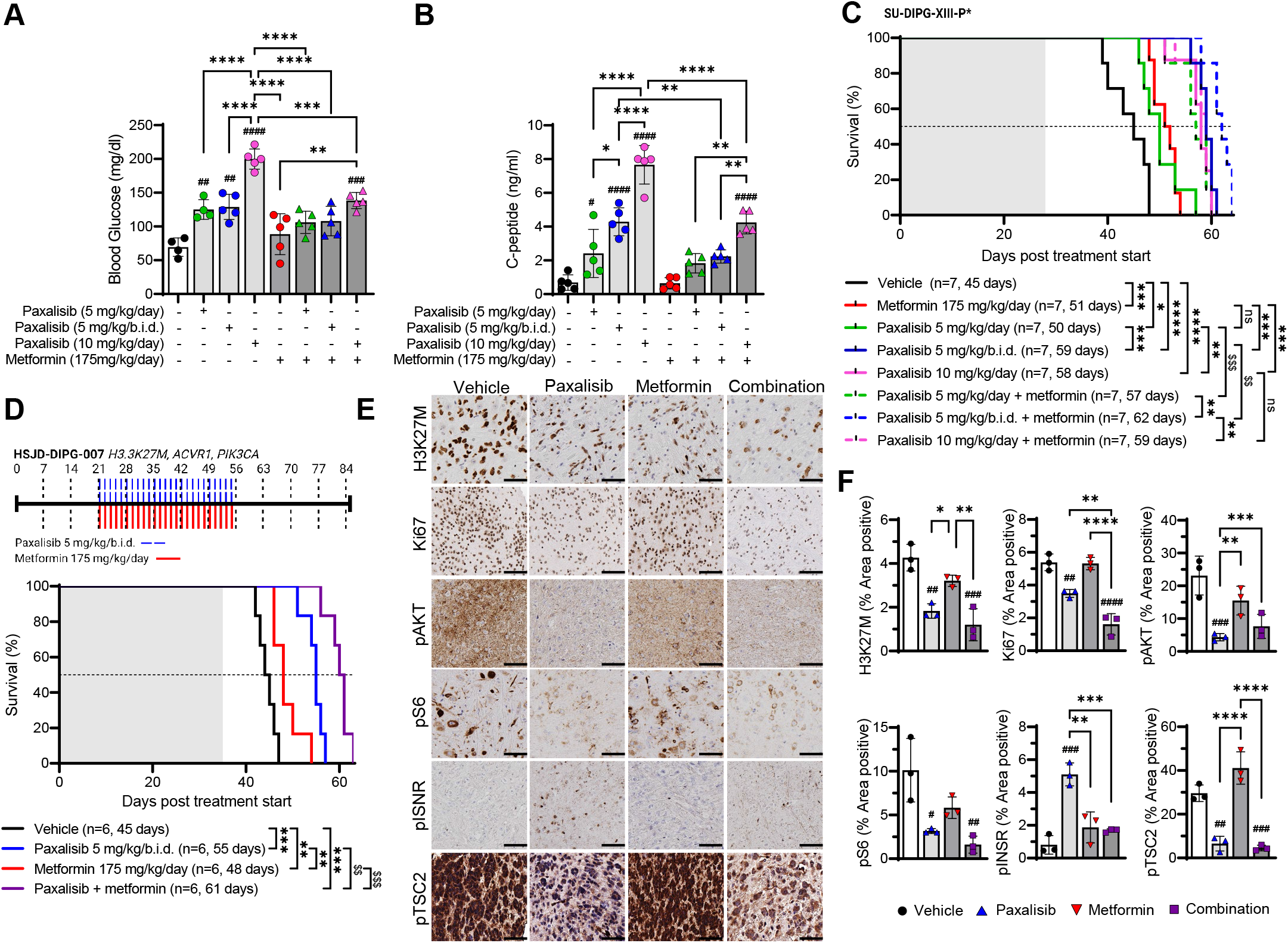
Efficacy of paxalisib is improved using metformin. (A) Blood glucose measurements 4 h post treatment with 5 mg/kg/day, 5 10 mg/kg/day paxalisib in combination with 175 mg/kg/day metformin. (B) C-peptide measurements 4 h post treatment with 5 mg/kg/b.i.d., 10 mg/kg/day paxalisib in combination with 175 mg/kg/day metformin. (C) Kaplan-Meier survival analysis for SU-xenografts treated with vehicle, paxalisib 5 mg/kg/day, 5 mg/kg/b.i.d., 10 mg/kg/day or each in combination with metformin 175 y gavage (analyzed by log rank test, ns=not significant, \**p<0.05*, \*\**p<0.01*, \*\*\**p<0.001*, \*\*\*\**p<0.0001*, synergistic comparisons*; p<0.001*). Shaded area indicates time receiving treatment. (D) Kaplan-Meier survival analysis for HSJD-DIPG-007 xenografts vehicle, paxalisib 5 mg/kg/b.i.d., metformin 175 mg/kg/day, or the combination of paxalisib and metformin by gavage (analyzed by, \**p<0.5*, \*\**p<0.01*, \*\*\**p<0.001*, \*\*\*\**p<0.0001*, synergistic comparisons; ^$$^*p<0.01*, ^$$$^*p<0.001*). Shaded area indicates time receiving) Tumor tissue resected from HSJD-DIPG-007 xenografts following 4 weeks of treatment and analyzed by Immunohistochemistry. e stained for H3K27M, pAKT, pS6, pTSC2 and pINSR (n=3 mice per treatment, representative images are presented, scale bar = Quantitation of IHC images using ImageJ (measured in technical triplicate, across biological replicates, n=3, one-way ANOVA, *<0.01*, \*\*\**p<0.001*, \*\*\*\**p<0.0001*, treated vs. untreated; ^#^*p<0.05*, ^##^*p<0.01*, ^###^*p<0.001*, ^####^*p<0.0001*).

To assess whether optimized paxalisib dosing improved survival when used alone and/or in combination with metformin, we employed the SU-DIPG-XIII-P* pontine orthotopic xenograft mouse model (Figure 4C). Encouragingly, 5 mg/kg/day, 5 mg/kg/b.i.d., 10 mg/kg/day paxalisib and metformin 175 mg/kg/day significantly increased survival compared to the vehicle controls. Furthermore, 5 mg/kg/b.i.d. and 10 mg/kg/day paxalisib significantly extended survival compared to 5 mg/kg/day. Metformin further potentiated the survival benefit of paxalisib 5 mg/kg/day and 5 mg/kg/b.i.d.; however, it did not provide additional benefit to the MTD 10 mg/kg/day regimen, highlighting that more frequent administration of a lower paxalisib dose in combination with metformin may provide the greatest clinical benefit for DMG patients. To further validate this finding, we examined the efficacy of paxalisib (5 mg/kg/b.i.d.) in combination with metformin in the HSJD-DIPG-007 (H3.3K27M, *PIK3CA-, ACVR1-*mutant) pontine orthotopic xenograft model. Supporting our previous finding, 5 mg/kg/b.i.d., paxalisib, again significantly extended survival compared to the vehicle (Figure 4D). The combination of paxalisib and metformin synergistically extended survival, compared to all treatments (Figure 4D). Immunohistochemistry showed that paxalisib treatment decreased phosphorylation of pAkt and pS6 *in vivo* (Figure 4E–F). However, using 5mg/kg/b.i.d. paxalisib alone, still increased phosphorylation of the insulin receptor (INSR) *in vivo* (Figure 4E–F), commensurate with the elevated C-peptide levels as seen in Figure 4B. The increased activity of the insulin pathway promoted by paxalisib treatment in DIPG xenograft mouse models was rescued using metformin (Figure 4E–F), promoting phosphorylation of tumor suppressor TSC2 at Thr1462 and reduced tumor burden as measured by H3K27M+ and Ki67 positive cells (Figure 4E–F).

### Paxalisib treatment promoted PKC signaling

To complement and extend the mechanistic insights established by RNA-seq (Figure 3), and garnish a view on the posttranslational landscapes of DIPG following paxalisib treatment, we performed global unbiased quantitative phosphoproteomic profiling of DIPG cells treated for 6 h with paxalisib (IC_50_ paxalisib, Figure 1B, Supplementary Table S6) A total of 6,017 unique proteins and 2,623 unique phosphoproteins were quantitatively identified across samples (Supplementary Figure S4A), with 753 significantly downregulated phosphoproteins and 95 significantly upregulated phosphoproteins (Supplementary Figure S4B). These analyses further confirmed paxalisib to be a potent inhibitor of the PI3K/Akt/mTOR pathway whilst simultaneously increasing phosphorylation of MARCKS and MARCKSL1, both substrates of active Protein kinase C (PKC) (Figure 5A). Kinases modulated in response to paxalisib were identified using Integrative inferred kinase activity (INKA) analysis (51) (Supplementary Figure S4C) and interrogated using IPA, which identified networks and canonical pathways mapping in GSK3B, mTOR and P70S6K, all regulated by PI3K (Figure 5B) and upstream regulators including AKT1, IGF1 and EGF (Figure 5C). PhoxTrack kinase activation analysis (52) identified pathways significantly upregulated by paxalisib treatment including CK2-A1, CSNK2A1, CK2, MAPKAPK2, PAK, and PKC signaling; these pathways either regulate intracellular calcium release or are influenced by calcium (Figure 5D). Interestingly, IPA analysis of RNA-seq data predicted a significant increase in IP_3_ signaling after paxalisib exposure (Figure 3A, Supplementary Figure S3B). IP_3_ is made through the hydrolysis of phosphatidylinositol 4,5-bisphosphate (PIP2), where it binds to its receptor (IP_3_R1) on the endoplasmic reticulum recycling and releasing calcium (Ca^2+^) into the cytoplasm (53).

**Figure 5:**
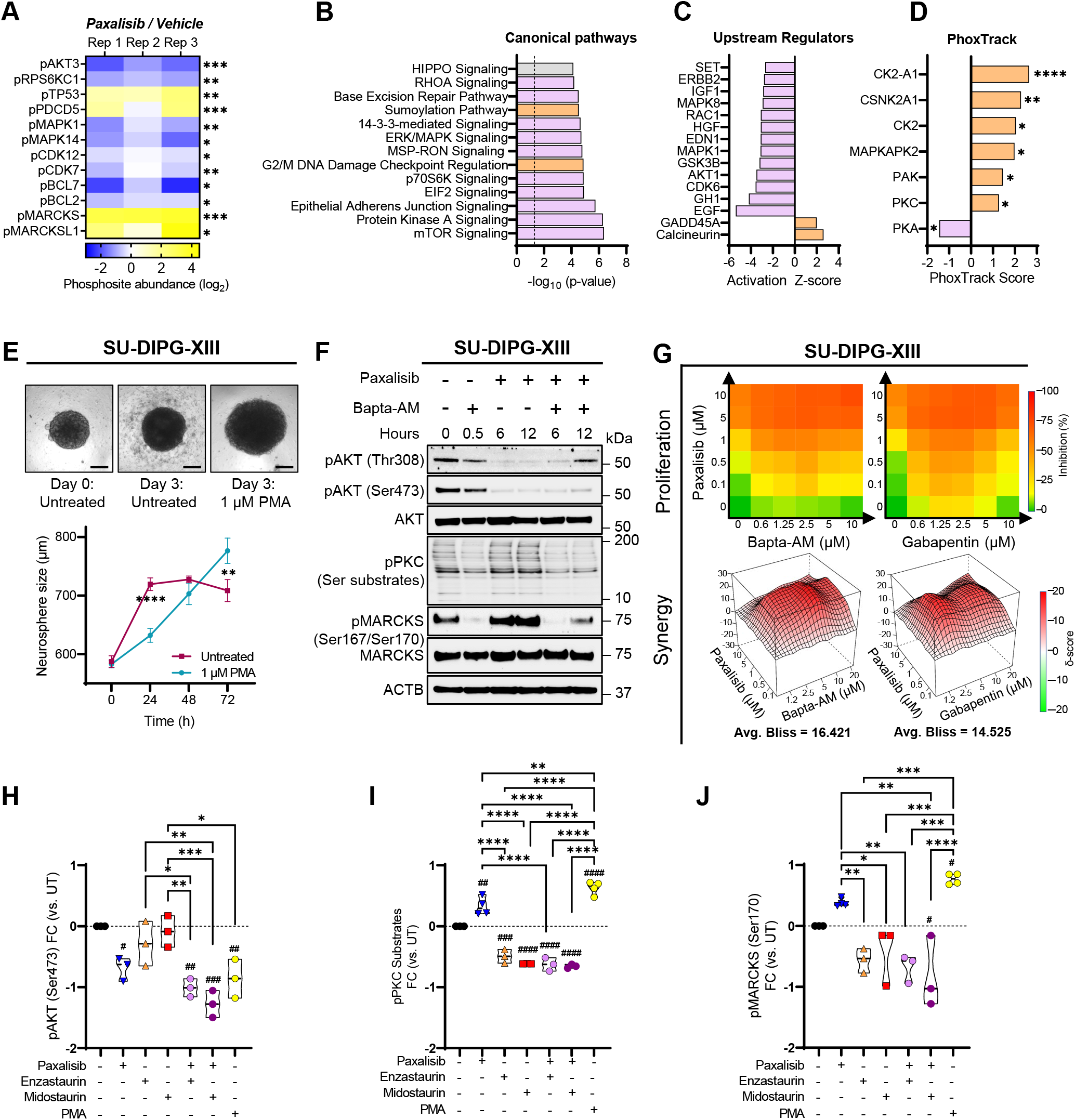
Phosphoproteomic analysis identified potent PKC activation following PI3K inhibition. (A) Significantly regulated oteins following paxalisib treatment (PI3K, MAPK, cell cycle, PKC and apoptotic signaling) (students t-test, \**p<0.05*, \*\**p<0.01*,). (B) Canonical pathways and (C) upstream regulators significantly altered by paxalisib exposure were identified by Ingenuity Analysis (IPA). Activated pathways have a positive z-score (purple), while inactivated pathways have a negative z-score (D) PhoxTrack analysis identified predicted kinases showing activation and inhibition following paxalisib treatment. Positive scores are shown in orange, while negative scores are in purple. (E) PKC was activated using Phorbol-12-myristate-13-acetate G-XIII neurospheres (scale bar 200 μM, two-way ANOVA, 1 µM PMA vs. Untreated \*\**p<0.01*, \*\*\*\**p<0.0001*). (F) BAPTA-AM was hibit paxalisib-induced PKC substrate and MARCKS phosphorylation measured by Western blotting (n=3, representative lots presented). (G) Bliss-synergy analysis of the combination of paxalisib with BAPTA-AM and Gabapentin. (H-J) Quantification g protein phosphorylation following combinations of paxalisib and PKC inhibitors after 24 h (n=3 biological replicates, one-way *p<0.05*, \*\**p<0.01*, \*\*\*\**p<0.0001*, treated vs. untreated; ^#^*p<0.05*, ^##^*p<0.01*, ^###^*p<0.001*, ^####^*p<0.0001*).

Calcium (Ca^2+^) plays fundamental and diversified roles in neuronal plasticity through the regulation of PKC signaling (54). Thus, we used Phorbol 12-myristate 13-acetate (PMA) as a PKC signaling activator to evaluate its effect in DIPG cell lines. Exogeneous activation of PKC using PMA significantly increased the growth of DIPG neurospheres compared to untreated controls (Figure 5E). Paxalisib treatment of DIPG models increased the phosphorylation of PKC substrates, as well as the phosphorylation of PKC substrate pMARCKS (Ser170), and this was ablated using the Ca^2+^ chelator BAPTA-AM (Figure 5F, Supplementary Figure S5), suggesting that Ca^2+^ promotes PKC signaling in response to PI3K inhibition in DIPG. Next, we assessed cytotoxicity using Ca^2+^ targeting compounds, including BAPTA-AM and the voltage-gated Ca^2+^ ion channel inhibitor, gabapentin. Both were found to be synergistic when combined with paxalisib (Figure 5G). Combining paxalisib with PKC inhibitors including enzastaurin and midostaurin, potently inhibited AKT signaling (Figure 5H, Supplementary Figure S6), and suppressed PKC substrate phosphorylation (Figure 5I) and MARCKS phosphorylation (Figure 5J, Supplementary Figure S6) previously promoted following PI3K/Akt inhibition (Figure 5A, I, J). Altogether, the use of Ca2+ chelators / channel blocks, or PKC inhibitors has the potential to suppress paxalisib-induced PKC activation and could be used as combination strategies to potentiate the anti-DIPG benefits of paxalisib.

### High-throughput drug screening assays confirms the preclinical utility of targets predicted by phosphoproteomic profiling

Monotherapeutic approaches to treat DMG have unequivocally failed patients (23, 24). To identify potential combination strategies, we performed high-throughput combination drug screening assays across a panel of DIPG neurosphere cell lines (n=9, including DIPGs harboring WT-H3, H3.1K27M, H3.3K27M and +/-PI3K-alterations) using paxalisib as a backbone, combined with clinically relevant compounds targeting the genes (Figure 3), and/or signaling pathways (Figure 5A–D) identified via RNA-seq and phosphoproteomic profiling, respectively. High level synergy with paxalisib was shown across DIPG lines when also targeting CDKs using ribociclib and palbociclib, EGFR and VEGFR using erlotinib and vandetanib, and PKC using midostaurin and enzastaurin (Figure 6A). To identify the best strategy to test in orthotopic DIPG preclinical models, we assessed the potential of each drug to penetrate the brain using CNS multi-parameter optimization (MPO) (55), correlating predicted CNS penetration, with synergy with paxalisib. These analyses identified erlotinib, vorinostat, ribociclib, enzastaurin, palbociclib and vandetanib as potential paxalisib combination strategies (Figure 6B).

**Figure 6:**
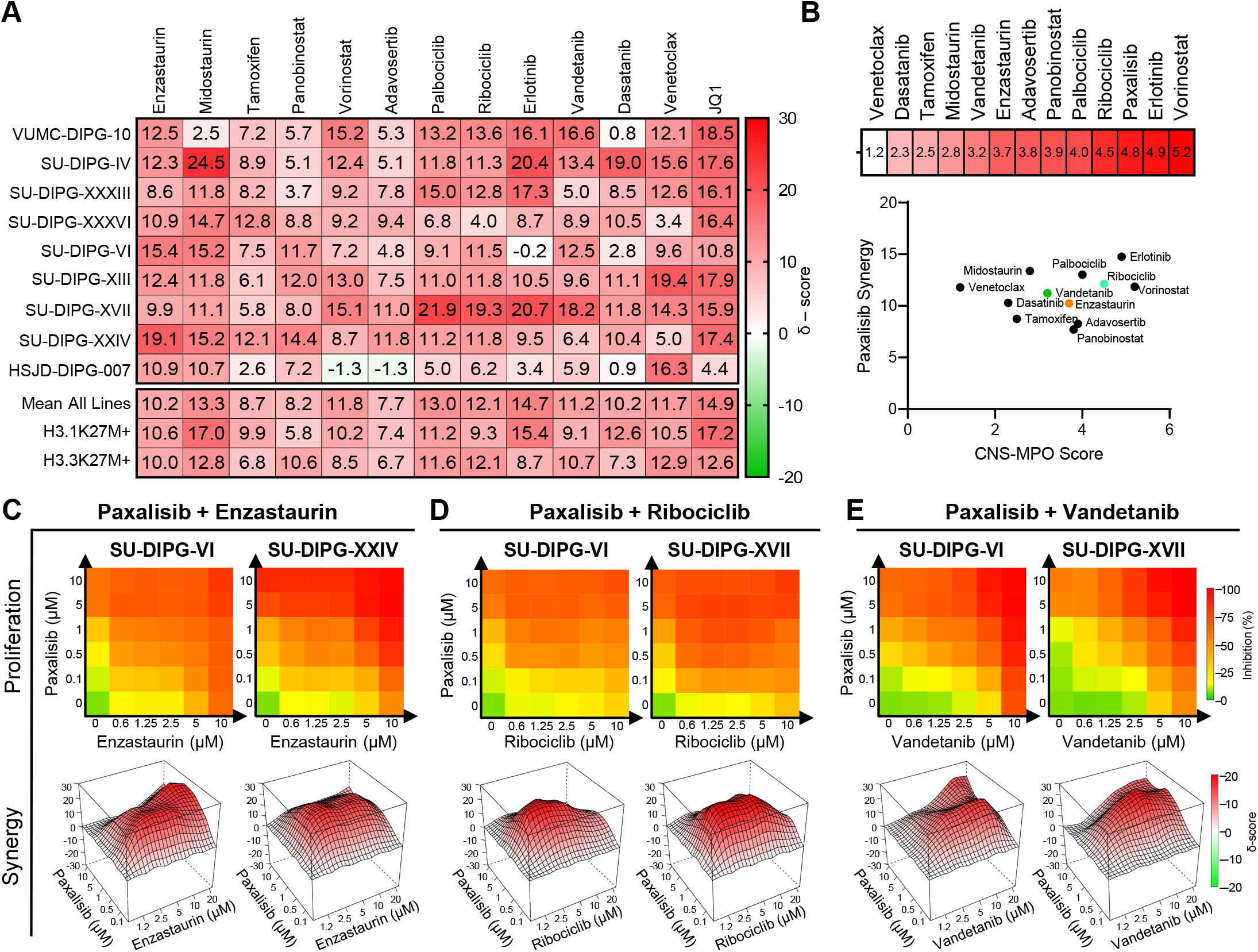
High-throughput drug screen identifies synergistic paxalisib drug combinations. (A) Bliss synergy analysis was performed using mbination with and clinically relevant inhibitors in a panel of DIPG cells lines (n=9), measured by resazurin cell growth and says after 72 h exposure (biological triplicate). (B) CNS-MPO analysis of was performed using compounds targeted pathways paxalisib treatment and plotted against their paxalisib combination synergy scores. Cell proliferation and bliss synergy analysis for n of (C) paxalisib and enzastaurin, (D) paxalisib and ribociclib and (E) paxalisib and vandetanib.

Enzastaurin (Figure 6C) (56), ribociclib (Figure 6D) (57) and vandetanib (Figure 6E) (58) all showed high-level synergism when combined with paxalisib across dose ranges, are brain penetrant, and FDA approved for other indications. Given that all these compounds have previously been tested in clinical trials as monotherapies for children diagnosed with DIPG and hence have known MTD and toxicity profiles, we therefore elected to test the preclinical efficacy of these compounds in combination with paxalisib using the aggressive DIPG orthotopic xenograft model SU-DIPG-XIII-P* (Figure 7). Enzastaurin (100 mg/kg/day), ribociclib (75 mg/kg/day), vandetanib (25 mg/kg/b.i.w.) and paxalisib (5 mg/kg/b.i.d.) all increased survival compared to vehicle controls (Figure 7A–C). However, only the combination of paxalisib and enzastaurin synergistically enhanced survival compared to both monotherapies (Figure 7A, Supplementary Figure S7A). The combination of paxalisib and ribociclib provided an additive survival benefit compared to both monotherapies (Figure 7B, Supplementary Figure S7B). The combination of paxalisib and vandetanib provided no additional survival benefit, potentially because of increased toxicity (observed significant weight loss) by administering both therapies orally, which necessitated a reduced treatment time of 2 weeks compared to 4 for the other combinations (Figure 7C, Supplementary Figure S7C). Given the encouraging paxalisib and enzastaurin combination xenograft results, we validated this efficacy using the DIPG model, RA055 (a biopsy derived post-radiation treatment model (59)), employing the optimized paxalisib dosing regimen (5 mg/kg/b.i.d., in combination with 175 mg/kg/day metformin, hereafter referred to as “optimized paxalisib”) which combined synergistically when compared to enzastaurin alone and provided an additive benefit compared to optimized paxalisib dosing (Figure 7D). Mice remained symptom free while on the combination of optimized paxalisib and enzastaurin; however, mice succumbed either due to neurological symptoms or weight loss, 9 days following the end of treatment (Figure 7D, Supplementary Figure S7D). Immunohistochemical analysis of tumors resected at the end of treatment (Figure 7E–F), mirrored the survival benefit, with optimized paxalisib and enzastaurin decreasing tumor burden as measured by H3K27M and Ki67, further decreased by combination treatment. Mechanistically, paxalisib decreased pAKT and subsequently increased PKC signaling commensurate with the *in vitro* results, and this was rescued by the treatment with enzastaurin in combination (Figure 7F).

**Figure 7:**
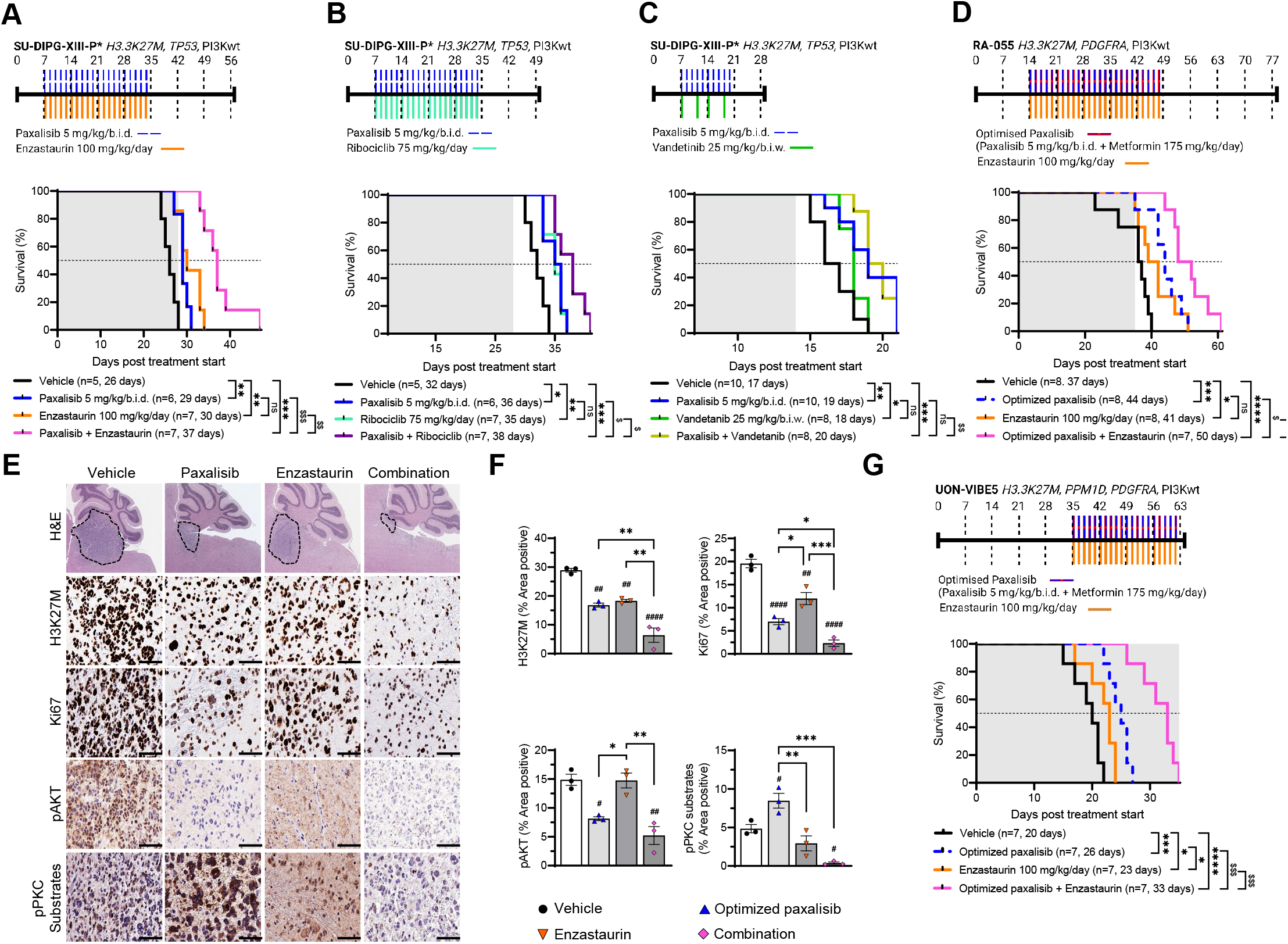
Synergistic combinations strategies targeting PI3K/Akt and PKC in DIPG xenograft mouse models. Kaplan-Meier survival analysis II-P* xenografts treated with paxalisib (5 mg/kg/b.i.d.) and compounds revealed to be synergistic by high through put drug screening uding (A) enzastaurin (100 mg/kg/day), (B) ribociclib (75 mg/kg/day) and (C) vandetanib (25 mg/kg/b.i.w.) (analyzed by log rank test, *.01*, \*\*\**p<0.001*, \*\*\*\**p<0.0001*, synergistic comparisons; ^$^*p<0.01*, ^$$^*p<0.01*, ^$$$^*p<0.001*). Shaded area indicates time receiving Kaplan-Meier survival analysis of RA-055 xenografts treated with the optimized combination of paxalisib (5 mg/kg/b.i.d.) + metformin y) and enzastaurin (100 mg/kg/day) (analyzed by log rank test, \**p<0.05*, \*\**p<0.01*, \*\*\**p<0.001*, \*\*\*\**p<0.0001, synergistic p<0.01*, ^$$^*p<0.01*, ^$$$^*p<0.001*). Shaded area indicates time receiving treatment. (E) Tumor tissue was resected from RA-055 wing 4 weeks of treatment (end of shaded area) and analyzed by immunohistochemistry. Sections were stained for H3K27M, Ki67, C Substrates (n=3 mice per treatment, representative images are presented, scale bar = 50 μM). (F) Quantitation of IHC images measured in technical triplicate, across biological replicates, n=3, one-way ANOVA, \**p<0.05*, \*\**p<0.01*, \*\*\**p<0.001*, \*\*\*\**p<0.0001*, reated; ^#^*p<0.05*, ^##^*p<0.01*, ^###^*p<0.001*, ^####^*p<0.0001*). (D) Kaplan-Meier survival analysis of UON-VIBE5 xenografts treated with the bination of paxalisib (5 mg/kg/b.i.d.) + metformin (175 mg/kg/day), and enzastaurin (100 mg/kg/day) (analyzed by log rank test, *01*, \*\*\**p<0.001*, \*\*\*\**p<0.0001*, synergistic comparisons; ^$^*p<0.01*, ^$$^*p<0.01*, ^$$$^*p<0.001*). Shaded area indicates time receiving

DIPG patients are often diagnosed with advanced disease through the clinical presentation of hallmark DIPG symptoms. To test if our optimized paxalisib and enzastaurin treatment regimen could provide a survival benefit to patients at disease progression or advanced disease stage, mice were xenografted with the DIPG patient derived autopsy cell line (UON-VIBE5, H3.3K27M, *PDGRFA*, *PPM1D*), employing a continuous treatment regimen commencing at the first sign of DIPG symptoms (weight loss). Here, vehicle treated mice succumbed 20 days post treatment start. Optimized paxalisib and enzastaurin provided survival benefits as monotherapies (26 days and 23 days, respectively); however, the combination of optimized paxalisib (Inclusive of metformin) and enzastaurin extended survival compared to both the vehicle and to each monotherapy (33 days), highlighting the preclinical potential of this combination for DMG patients at either the upfront, or advanced disease setting settings.

## Discussion

DIPG is an insidious disease responsible for more deaths in children than any other cancer (10, 23, 60). Hallmark hypomethylated euchromatic landscapes instigated by H3-alterations (5) drive transcriptional programs that promote cellular immortality (61). Transcriptional volatility, coupled with cooccurring somatic mutations in tumor suppressor and signaling genes (3), offers some explanation as to why standard-of-care radiotherapy provides only a transient benefit, and monotherapeutic treatment strategies have failed DIPG patients (10, 23, 24). The PI3K/Akt/mTOR signaling cascade lies immediately downstream of many upregulated and mutant growth factor receptors responsible for the transmission of oncogenic signals that promote proliferation, angiogenesis and metabolism, making the pathway an attractive therapeutic target. The importance of PI3K-mTOR signaling uncovered herein by analysis of loss-of-function CRISPR-Cas9 screen data, reveals both the catalytic p110α subunit of PI3K (*PIK3CA*) and downstream serine/threonine protein kinase *MTOR* are required to sustain DIPG cell growth and proliferation *in vitro*. Given the recurring nature of PI3K-related gene mutations compared to mTOR in DIPG, we focused these studies on testing and optimizing the brain penetrant PI3K-inhibitor paxalisib, optimized to penetrate the CNS and shown to disperse throughout the brain (17), and in particular in the brainstem following the dose optimization regimen identified herein (Figure 2D).

Activated PI3K/Akt signaling, particularly through PIK3CA and RAC-beta serine/threonine-protein kinase (AKT2) mediate insulin-driven glucose uptake in muscle, liver, and fat cells, following translocation of glucose transporters to the plasma membrane (62). Hence, PI3K/Akt inhibition blocks insulin-driven glucose uptake resulting in a dose-dependent increase in plasma levels of fasting C-peptide and insulin, to cause hyperglycemia (63). Insulin is a systemic obligatory on-target pharmacodynamic surrogate for PI3K inhibition, activating the insulin receptor and reactivating PI3K/Akt signaling, particularly in DIPG characterized by an abundance of insulin receptors (64, 65), potentially limiting the clinical benefit of PI3K antagonists (62).

PI3K/Akt pathway inhibitors commonly cause toxicities that are dose limiting. Phase 1b clinical trials testing safety, tolerability, and pharmacokinetics and to estimate the MTD of paxalisib when administered immediately post-radiotherapy in the pediatric DIPG setting (NCT03696355), identified DLTs including hyperglycemia and mucositis (18). This clinical trial established a safe dose of 27 mg/m^2^/day, equating to an equivalent mouse dose of 9.2 mg/kg/day (50). Here, mice treated with 10 mg/kg/day showed significantly elevated blood glucose and C-peptide levels indicative of hyperglycemia and hyperinsulinemia. By contrast, an optimized 5 mg/kg/b.i.d. dosing regimen decreased blood glucose levels below that recognized as hyperglycemic but still elevated compared to vehicle control treated mice. Such results raise the prospect of exploiting treatment paradigms that combine the use of anti-glycemic approaches, such as metformin, in tandem with PI3K inhibitors to maintain glucose homeostasis like trials testing paxalisib plus metformin, or paxalisib plus a ketogenic diet in the adult brain cancer setting (NCT05183204).

The improved twice daily 5 mg/kg regimen we adopted herein increased the survival of mice compared to once daily 10 mg/kg, suggestive of an accumulation of paxalisib in the brainstem and sustained inhibition of PI3K/Akt signaling (Figure 4C, D). This benefit was potentiated using systemic control of insulin via metformin administration; however, this response was restricted to the use of either 5 mg/kg/day or 5 mg/kg/b.i.d. paxalisib. Metformin’s primary target is the liver, where it decreases hepatic gluconeogenesis and stimulates glucose uptake in muscle. Like the liver, treatment of DIPG orthotopic xenograft mouse models promoted AMPK independent phosphorylation and activation of the mTOR tumor suppressor TSC2 (66) to decrease mTOR activity and protein synthesis to provide additional control over DIPG cell growth and survival (Figure 4E, F). Previous PK studies in the brain of mice showed metformin to be active in the CNS and to reach midline structures at a plasma to brain ratio of 1:0.99 (67).

Studies have shown *in vitro* anti-DIPG benefits from combining metformin and the pyruvate dehydrogenase kinase inhibitor dichloroacetate, a strategy that decreased proliferation and caused apoptosis; however, in this situation metformin did not significantly improve the survival of DIPG xenograft mice when used alone at 125 mg/kg/b.i.d. (68). We initially performed MTD studies using the combination of paxalisib 5 mg/kg/b.i.d., and metformin using 250 mg/kg/day in line with studies that showed this dose achieved potent mTOR antagonism in vivo (66). However, mice lost weight rapidly; therefore, we treated mice with metformin at 175 mg/kg/day, alone and in combination with paxalisib 5 mg/kg/b.i.d. This dose of metformin equates to ∼14 mg/kg/day or (840 mg/60kg adult) analogous to clinical dosing for patients with type 2 diabetes (69). The combination of metformin and paxalisib extended the survival of mice synergistically compared to both monotherapies, thus serving as an important consideration for ongoing clinical studies testing paxalisib.

DIPGs display a high degree of intra-tumoral clonal diversity (9, 23, 70), highlighting the necessity to develop effective combination strategies to improve survival. Focal gains in *PDGFRA*, *EGFR* and *VEGFR* are seen in ∼32% of DIPG cases, with PI3K alterations including constitutive activating mutations in *PIK3CA*, *PIK3R1* and loss of function of *PTEN* (seen in 43% of patients combined), the latter associated with worse overall survival in DIPG (71), all of which promote constitutive PI3K/Akt/mTOR signaling (10). The novel application of RNA-sequencing and phosphoproteomic profiling of DIPG cells treated with paxalisib identified increased Ca^2+^-activated PKC signaling. PRKCB (PKCβ) is a serine/threonine-protein kinase involved in various cellular processes including insulin signaling, energy metabolism, and regulation of the B-cell receptor (BCR) signalosome (72). PKC is also activated following the binding of Brain-derived neurotrophic factor (BDNF) to Neurotrophic receptor tyrosine kinase 2 (NTRK2, or TrkB), opening AMPAR channels to the postsynaptic membrane, again fundamentally regulated by Ca^2+^ (73). These processes are not only critical in the control of learning and behavior, but underpin neuron-DIPG and DIPG-DIPG communications (73). This encouraged us to test the CNS active PKC inhibitor enzastaurin in combination with paxalisib which led to synergistic survival extension of DIPG xenograft models, highlighting the promise of this novel combination approach. Our *in vitro* data further suggest that the voltage-gated Ca^2+^ ion channel inhibitor, gabapentin acts synergistically with paxalisib to decrease the growth and proliferation of DIPG cell lines (Figure 5G). Future studies to inhibit the role Ca^2+^ plays in the transmission of oncogenic signals in DIPG, in combination with optimized PI3K/Akt targeting might therefore go some way toward improving response for patients using these combinations. Hyperglycemia is a key factor responsible for the development of diabetic vascular complications through activation of PKC signaling (74). PKC propagates transmission of signals downstream of the insulin receptor, through a dose-and time-dependent increase in the phosphorylation of MARCKS (75) analogous to DIPG cells treated with paxalisib. Therefore, the use of anti-glycemic therapies in combination with PI3K and PKC inhibitors may increase the DIPG specific response. For example, metformin may further potentiate the therapeutic benefits of this multi-agent, anti-DIPG strategy. Encouragingly, xenograft mice treated with paxalisib and metformin in combination with the PKCβ inhibitor enzastaurin, remained symptom free while on therapy, highlighting the rational inclusion of clinically relevant therapies targeting the emerging biology revealed by our multiomic drug profiling strategy (Figures 3 and 5).

The preclinical investigation described herein has optimized paxalisib for the treatment of DIPG. Here, we have potentiated the on-target efficacy of the drug, whilst reducing common PI3K/Akt inhibitor related side-effects and toxicities. Further, by performing multiomic analysis of DIPG models treated with paxalisib we have identified combination strategies that are targetable using FDA approved therapies. Our data supports a model by which PI3K inhibition in DIPG, drives calcium dependent PKC activation and in the process uncovers a clinically actionable combination strategy to inform future clinical trials for patients.

## Acknowledgments

We acknowledge all children and their families diagnosed with DIPG/DMG. We are sincerely grateful to all families who donated tissue in support of our DIPG/DMG research. We thank Dr Michelle Monje, Stanford University, CA; Dr Esther Hulleman, Princess Máxima Center for Pediatric Oncology, Netherlands, Dr Angel M. Carcaboso, Department of Pediatric Hematology and Oncology, Spain, for the generous donation of DIPG neurosphere cell lines used in this study. The UON-JUMP4 cell line was established at The University of Newcastle with thanks to the Zero Childhood Cancer Program, Children’s Cancer Institute, Australia. UON-VIBE5 cell line was established at The University of Newcastle by Dr Ryan Duchatel and A/Prof Matthew Dun. Proteomics was supported by Nathan Smith from The Analytical and Biomolecular Research Facility and The Academic and Research Computing Support (ARCS) team, within IT Services at the University of Newcastle, who provided high performance computing (HPC) infrastructure for supporting the bioinformatics. Histology services were provided by Cassandra Griffin, Fiona Richards, Megan Clarke, and Kaylee O’Brien from the HMRI Core Histology Facility. Immunohistochemistry optimization and staining services were provided by The University of Newcastle’s ‘NSW Regional Biospecimen Services’, with support from NSW Health Pathology. Figures were created with the assistance of BioRender.com.

## Supplementary Data Captions

**Supplementary Figure S1: Expression and phosphorylation of PI3K/Akt/mTOR signaling proteins does not predict sensitivity to with paxalisib in vitro**.

(A) Growth rates of SU-DIPG-XXXVI and SU-DIPG-XIII cells harboring *PIK3CA* knockout. (B) Protein expression of key PI3K/Akt/mTOR signaling proteins in DIPG cell lines measured by Western blot (n=3, representative Western blots presented). (C) Densitometry of Western blot values normalized to VUMC10 and correlated to IC25 values for paxalisib (significance determined using Pearson’s linear regression). (D) Protein phosphorylation of key PI3K/Akt/mTOR signaling proteins in DIPG cell lines (SU-DIPG-VI, SU-DIPG-XIII, SU-DIPG-XVII) treated with paxalisib over time (1 µM, n=3, representative Western blots presented).

**Supplementary Figure S2: Paxalisib dose optimization maintains inhibition of PI3K/Akt/mTOR signaling.** (A) SU-DIPG-XIII-P* bearing xenograft mice weights were measure after treated with either vehicle, 5 mg/kg/day, 5 mg/kg/b.i.d., or 10 mg/kg/day treatment regime over time. (B) Tumors were resected from mice treated with vehicle, 6, 12 and 24 h, post treatment after 2 weeks of paxalisib treatment (vehicle, 5mg/kg/day, 5mg/kg/b.i.d., or 10mg/kg/day) and phosphorylation and expression of key PI3K/Akt/mTOR related signaling proteins measured by Western blot (n=3, representative Western blots presented).

**Supplementary Figure S3: Volcano plot analysis of transcription regulation following paxalisib treatment**.

(A) Bulk RNA Barcoding and sequencing (BRB-seq) of SU-DIPG-VI following paxalisib treatment (1 µM) for pooled data. (B) Validation of important oncogenes shown to be significantly modulated by paxalisib treatment at the protein level by Western blot (n=3, representative Western blots presented).

**Supplementary Figure S4: Phosphoproteomic profiling of paxalisib treated DIPG models**.

(A) Phosphoproteomic profiling of SU-DIPG-XXXVI cells treated with 1 µM paxalisib for 6 h (n=3, biological triplicate). (B) Volcano plot of differentially phosphorylated proteins, (p<0.05 log2 fold change <-0.6 and >0.6, n=3). (C) Integrated kinase expression analysis using Integrative inferred kinase activity (INKA) analysis of paxalisib treated cells (n=3).

**Supplementary Figure S5: Increased calcium dependent PKC following paxalisib treatment**.

Quantification of Western blot measurement of SU-DIPG-XIII cells treated with paxalisib or the calcium chelator BAPTA-AM (A) pAKT (Thr308), (B) pAKT(Ser473), (C) pMARCKS (Ser170) and (D) pSerPKC Substrates (n=3, one-way ANOVA, *\*p<0.05, **p<0.01, ***p<0.001, ****p<0.0001*).

**Supplementary Figure S6: Assessment of PI3K-and PKC-related signaling following the combination of paxalisib with PKC inhibitors**.

Protein expression and phosphorylation of key PI3K/Akt/mTOR and PKC signaling proteins was assessed after exposure to paxalisib (1 µM, 24 h), enzastaurin (5 µM, 24 h), midostaurin (5 µM, 24 h) and the PKC activator, PMA (1 µM, 24 h) (n=3, biological replicates, representative Western blots presented).

**Supplementary Figure S7: DIPG xenograft mouse weight following treatment with paxalisib alone and in combination with FDA approved therapies predicted to synergize by phosphoproteomic profiling**.

Mouse weights recorded for SU-DIPG-XIII-P* xenograft models, following treatment with vehicle, paxalisib (5 mg/kg/b.i.d.) alone and combined with (A) enzastaurin (100 mg/kg/day), (B) ribociclib (75 mg/kg/day) or (C) vandetanib (25 mg/kg/b.i.w.). (D) RA-055 xenograft mouse model weights, following treatment with vehicle, optimized paxalisib (5 mg/kg/b.i.d., + 175 mg/kg/day metformin) in combination with enzastaurin (100 mg/kg/day).

## Supplementary Tables

Supplementary Table S1: Primary antibodies used for Western immunoblotting and immunohistochemistry.

Supplementary Table S2: Cell line sensitivity to paxalisib.

Supplementary Table S3: Differential gene expression data for DIPG cells treated for 6 h with paxalisib.

Supplementary Table S4: Differential gene expression data for DIPG cells treated for 12 h with paxalisib.

Supplementary Table S5: Pooled differential gene expression data for DIPG cells treated for 6 and 12 h with paxalisib.

Supplementary Table S6: Significantly regulated phosphorylation sites for DIPG cells treated for 6 h with paxalisib.

## Notes

**Funding**: R.J.D., is supported by a ChadTough DefeatDIPG Foundation Postdoctoral Fellowship. M.D.D., was supported by a Cancer Institute NSW Fellowship. M.D.D., holds an NHMRC Investigator Grant – GNT1173892. The contents of the published material are solely the responsibility of the research institutions involved or individual authors and do not reflect the views of NHMRC. M.D.D., was also supported by a ChadTough DefeatDIPG New Investigator Fellowship. J.E.C., is supported by a Victorian Cancer Agency Mid-Career Fellowship (MCRF17014). The work at the Hudson was supported by the Children’s Cancer Foundation and an MRFF Grant (APP2007620). E.R.J., is supported by the Hunter Cancer Research Alliance and the Josephine Dun Scholarship from the Isabella and Marcus Foundation and Miette Skiller Scholarship Fund a sub-fund of the Australian Communities Foundation. S.G.P., is supported by the Hudson Institute of Medical Research and the Children’s Cancer Foundation. I.J.F., is recipient of the RUN DIPG ‘Moving Towards a Cure’ HDR Scholarship and CureCell ExCELLerate Award. M.L.P., is recipient of the RUN DIPG International HDR Scholarship. This project (COMBATT DMG 1.0 and 2.0) was supported by the DIPG/DMG Collaborative which is supported by The Cure Starts Now Foundation, The Cure Starts Now Foundation Australia, Brooke Healey Foundation, Wayland Villars Foundation, ChadTough Foundation, Aidan’s Avengers, Austin Strong, Cure Brain Cancer, Jeffrey Thomas Hayden Foundation, Laurie’s Love Foundation, Love Chloe Foundation, Musella Foundation, Pray Hope Believe, Reflections Of Grace, Storm the Heavens Fund, Aubreigh’s Army, Whitley’s Wishes, Ryan’s Hope, Benny’s World, The Isabella and Marcus Foundation, Lauren’s Fight for Cure, Robert Connor Dawes Foundation, The Gold Hope Project, Julia Barbara Foundation, Lily Larue Foundation, American Childhood Cancer Organization, RUN DIPG, Gabriella’s Smile Foundation, and Snapgrant.com. Funding also received from Strategic Group, McDonald Jones Foundation, Vinva Foundation, Kiriwina Investments, Keith Tulloch Wines, Pacific Pediatric Neuro-Oncology Consortium Foundation, Yuvaan Tiwari Foundation, Edie’s Kindness Project, Maitland Cancer Appeal, Blackjack Foundation, and The Hunter Medical Research Institute. Project funding was provided by Kazia Therapeutics. The Cancer Institute NSW and NMHRC in partnership with the College of Health, Medicine and Wellbeing at the University of Newcastle funded the mass spectroscopy platform. D.S.Z. and M.T. are supported by a Cancer Institute NSW Translational Research Program Grant. H.C.M is supported by a Cancer Institute NSW Fellowship (ECF1299).

**Conflicts of Interest**: M.D.D. is a parent to a child lost to diffuse intrinsic pontine glioma (DIPG), and the Founder and a Director of the not-for-profit charity RUN DIPG Pty Ltd. D.S.Z reports consulting / advisory board fees from Bayer, Astra Zeneca, Accendatech, Novartis, Day One, FivePhusion, Amgen, Alexion, and Norgine and research support from Accendatech. Kazia Therapeutics contributed project funding supporting the animal studies outlined herein. Paxalisib was provided by Kazia Therapeutics under material transfer agreement (MTA). Enzastaurin was provided by Denovo Biopharma under MTA.

### Competing Interest Statement

M.D.D. is a parent to a child lost to diffuse intrinsic pontine glioma (DIPG), and the Founder and a Director of the not-for-profit charity RUN DIPG Pty Ltd. D.S.Z reports consulting / advisory board fees from Bayer, Astra Zeneca, Accendatech, Novartis, Day One, FivePhusion, Amgen, Alexion, and Norgine and research support from Accendatech. Kazia Therapeutics contributed project funding supporting the animal studies outlined herein. Paxalisib was provided by Kazia Therapeutics under material transfer agreement (MTA). Enzastaurin was provided by Denovo Biopharma under MTA.

## References

1. Vanan MI, and Eisenstat DD. DIPG in Children – What Can We Learn from the Past? Front Oncol. 2015;5:237.

2. Cooney T, Lane A, Bartels U, Bouffet E, Goldman S, Leary SES, et al. Contemporary survival endpoints: an International Diffuse Intrinsic Pontine Glioma Registry study. Neuro Oncol. 2017;19(9):1279–80.

3. Mackay A, Burford A, Carvalho D, Izquierdo E, Fazal-Salom J, Taylor KR, et al. Integrated Molecular Meta-Analysis of 1,000 Pediatric High-Grade and Diffuse Intrinsic Pontine Glioma. Cancer Cell. 2017;32(4):520–37 e5.

4. Bender S, Tang Y, Lindroth AM, Hovestadt V, Jones DT, Kool M, et al. Reduced H3K27me3 and DNA hypomethylation are major drivers of gene expression in K27M mutant pediatric high-grade gliomas. Cancer Cell. 2013;24(5):660–72.

5. Lewis PW, Muller MM, Koletsky MS, Cordero F, Lin S, Banaszynski LA, et al. Inhibition of PRC2 activity by a gain-of-function H3 mutation found in pediatric glioblastoma. Science. 2013;340(6134):857–61.

6. Chan KM, Fang D, Gan H, Hashizume R, Yu C, Schroeder M, et al. The histone H3.3K27M mutation in pediatric glioma reprograms H3K27 methylation and gene expression. Genes Dev. 2013;27(9):985–90.

7. Hubner JM, Muller T, Papageorgiou DN, Mauermann M, Krijgsveld J, Russell RB, et al. EZHIP/CXorf67 mimics K27M mutated oncohistones and functions as an intrinsic inhibitor of PRC2 function in aggressive posterior fossa ependymoma. Neuro Oncol. 2019;21(7):878–89.

8. Castel D, Kergrohen T, Tauziède-Espariat A, Mackay A, Ghermaoui S, Lechapt E, et al. Histone H3 wild-type DIPG/DMG overexpressing EZHIP extend the spectrum diffuse midline gliomas with PRC2 inhibition beyond H3-K27M mutation. Acta Neuropathologica. 2020;139(6):1109–13.

9. Nikbakht H, Panditharatna E, Mikael LG, Li R, Gayden T, Osmond M, et al. Spatial and temporal homogeneity of driver mutations in diffuse intrinsic pontine glioma. Nat Commun. 2016;7:11185.

10. Duchatel RJ, Jackson ER, Alvaro F, Nixon B, Hondermarck H, and Dun MD. Signal Transduction in Diffuse Intrinsic Pontine Glioma. Proteomics. 2019;19(21–22):e1800479.

11. Lien EC, Dibble CC, and Toker A. PI3K signaling in cancer: beyond AKT. Curr Opin Cell Biol. 2017;45:62–71.

12. Yang J, Nie J, Ma X, Wei Y, Peng Y, and Wei X. Targeting PI3K in cancer: mechanisms and advances in clinical trials. Mol Cancer. 2019;18(1):26.

13. Hess G, Herbrecht R, Romaguera J, Verhoef G, Crump M, Gisselbrecht C, et al. Phase III study to evaluate temsirolimus compared with investigator’s choice therapy for the treatment of relapsed or refractory mantle cell lymphoma. J Clin Oncol. 2009;27(23):3822–9.

14. Pavel ME, Hainsworth JD, Baudin E, Peeters M, Horsch D, Winkler RE, et al. Everolimus plus octreotide long-acting repeatable for the treatment of advanced neuroendocrine tumours associated with carcinoid syndrome (RADIANT-2): a randomised, placebo-controlled, phase 3 study. Lancet. 2011;378(9808):2005-12.

15. Furman RR, Sharman JP, Coutre SE, Cheson BD, Pagel JM, Hillmen P, et al. Idelalisib and rituximab in relapsed chronic lymphocytic leukemia. N Engl J Med. 2014;370(11):997–1007.

16. Brennan CW, Verhaak RG, McKenna A, Campos B, Noushmehr H, Salama SR, et al. The somatic genomic landscape of glioblastoma. Cell. 2013;155(2):462–77.

17. Salphati L, Alicke B, Heffron TP, Shahidi-Latham S, Nishimura M, Cao T, et al. Brain Distribution and Efficacy of the Brain Penetrant PI3K Inhibitor GDC-0084 in Orthotopic Mouse Models of Human Glioblastoma. Drug Metab Dispos. 2016;44(12):1881–9.

18. Tinkle C, Huang J, Campagne O, Pan H, Onar-Thomas A, Chiang J, et al. CTNI-27. FIRST-IN-PEDIATRICS PHASE I STUDY OF GDC-0084 (PAXALISIB), A CNS-PENETRANT PI3K/mTOR INHIBITOR, IN NEWLY DIAGNOSED DIFFUSE INTRINSIC PONTINE GLIOMA (DIPG) OR OTHER DIFFUSE MIDLINE GLIOMA (DMG). Neuro-Oncology. 2020;22(Suppl 2): ii48.

19. Wen PY, Cloughesy TF, Olivero AG, Morrissey KM, Wilson TR, Lu X, et al. First-in-Human Phase I Study to Evaluate the Brain-Penetrant PI3K/mTOR Inhibitor GDC-0084 in Patients with Progressive or Recurrent High-Grade Glioma. Clin Cancer Res. 2020;26(8):1820–8.

20. Hopkins BD, Pauli C, Du X, Wang DG, Li X, Wu D, et al. Suppression of insulin feedback enhances the efficacy of PI3K inhibitors. Nature. 2018;560(7719):499-503.

21. Ippen FM, Alvarez-Breckenridge CA, Kuter BM, Fink AL, Bihun IV, Lastrapes M, et al. The Dual PI3K/mTOR Pathway Inhibitor GDC-0084 Achieves Antitumor Activity in PIK3CA-Mutant Breast Cancer Brain Metastases. Clin Cancer Res. 2019;25(11):3374–83.

22. Mishra R, Patel H, Alanazi S, Kilroy MK, and Garrett JT. PI3K Inhibitors in Cancer: Clinical Implications and Adverse Effects. Int J Mol Sci. 2021;22(7).

23. Findlay IJ, De Iuliis GN, Duchatel RJ, Jackson ER, Vitanza NA, Cain JE, et al. Pharmaco-proteogenomic profiling of pediatric diffuse midline glioma to inform future treatment strategies. Oncogene. 2022;41(4):461–75.

24. André N, Buyens G. E. B. D. W, and Dun MD. Access to new drugs in paediatric oncology: can we learn from the ongoing ONC201 saga? The Lancet Oncology. 2023.

25. Sun C, Daniel P, Bradshaw G, Shi H, Loi M, Chew N, et al. Generation and Multi-Dimensional Profiling of a Childhood Cancer Cell Line Atlas Defines New Therapeutic Opportunities. Cancer Cell. 2023.

26. Murray HC, Enjeti AK, Kahl RGS, Flanagan HM, Sillar J, Skerrett-Byrne DA, et al. Quantitative phosphoproteomics uncovers synergy between DNA-PK and FLT3 inhibitors in acute myeloid leukaemia. Leukemia. 2020.

27. Dun MD, Anderson AL, Bromfield EG, Asquith KL, Emmett B, McLaughlin EA, et al. Investigation of the expression and functional significance of the novel mouse sperm protein, a disintegrin and metalloprotease with thrombospondin type 1 motifs number 10 (ADAMTS10). Int J Androl. 2012;35(4):572–89.

28. Pestinger V, Smith M, Sillo T, Findlay JM, Laes J-F, Martin G, et al. Use of an Integrated Pan-Cancer Oncology Enrichment Next-Generation Sequencing Assay to Measure Tumour Mutational Burden and Detect Clinically Actionable Variants. Molecular Diagnosis & Therapy. 2020;24(3):339–49.

29. Pan B, Kusko R, Xiao W, Zheng Y, Liu Z, Xiao C, et al. Similarities and differences between variants called with human reference genome HG19 or HG38. BMC Bioinformatics. 2019;20(2):101.

30. Dunn T, Berry G, Emig-Agius D, Jiang Y, Lei S, Iyer A, et al. Pisces: an accurate and versatile variant caller for somatic and germline next-generation sequencing data. Bioinformatics. 2019;35(9):1579–81.

31. Paila U, Chapman BA, Kirchner R, and Quinlan AR. GEMINI: integrative exploration of genetic variation and genome annotations. PLoS Comput Biol. 2013;9(7):e1003153.

32. Stodolna A, He M, Vasipalli M, Kingsbury Z, Becq J, Stockton JD, et al. Clinical-grade whole-genome sequencing and 3’ transcriptome analysis of colorectal cancer patients. Genome Med. 2021;13(1):33.

33. Chen X, Schulz-Trieglaff O, Shaw R, Barnes B, Schlesinger F, Källberg M, et al. Manta: rapid detection of structural variants and indels for germline and cancer sequencing applications. Bioinformatics. 2016;32(8):1220–2.

34. McLaren W, Gil L, Hunt SE, Riat HS, Ritchie GR, Thormann A, et al. The Ensembl Variant Effect Predictor. Genome Biol. 2016;17(1):122.

35. Magi A, Benelli M, Yoon S, Roviello F, and Torricelli F. Detecting common copy number variants in high-throughput sequencing data by using JointSLM algorithm. Nucleic Acids Res. 2011;39(10):e65.

36. Gu Z, Eils R, and Schlesner M. Complex heatmaps reveal patterns and correlations in multidimensional genomic data. Bioinformatics. 2016;32(18):2847–9.

37. Przystal JM, Cianciolo Cosentino C, Yadavilli S, Zhang J, Laternser S, Bonner ER, et al. Imipridones affect tumor bioenergetics and promote cell lineage differentiation in diffuse midline gliomas. Neuro Oncol. 2022;24(9):1438–51.

38. Alpern D, Gardeux V, Russeil J, Mangeat B, Meireles-Filho ACA, Breysse R, et al. BRB-seq: ultra-affordable high-throughput transcriptomics enabled by bulk RNA barcoding and sequencing. Genome Biol. 2019;20(1):71.

39. Schrode N, Seah C, Deans PJM, Hoffman G, and Brennand KJ. Analysis framework and experimental design for evaluating synergy-driving gene expression. Nat Protoc. 2021;16(2):812–40.

40. Love MI, Huber W, and Anders S. Moderated estimation of fold change and dispersion for RNA-seq data with DESeq2. Genome Biol. 2014;15(12):550.

41. Dun MD, Chalkley RJ, Faulkner S, Keene S, Avery-Kiejda KA, Scott RJ, et al. Proteotranscriptomic Profiling of 231-BR Breast Cancer Cells: Identification of Potential Biomarkers and Therapeutic Targets for Brain Metastasis. Mol Cell Proteomics. 2015;14(9):2316–30.

42. Degryse S, de Bock CE, Demeyer S, Govaerts I, Bornschein S, Verbeke D, et al. Mutant JAK3 phosphoproteomic profiling predicts synergism between JAK3 inhibitors and MEK/BCL2 inhibitors for the treatment of T-cell acute lymphoblastic leukemia. Leukemia. 2018;32(3):788–800.

43. Nixon B, Johnston SD, Skerrett-Byrne DA, Anderson AL, Stanger SJ, Bromfield EG, et al. Modification of Crocodile Spermatozoa Refutes the Tenet That Post-testicular Sperm Maturation Is Restricted To Mammals. Mol Cell Proteomics. 2019;18(Suppl 1):S58–S76.

44. Staudt DE, Murray HC, Skerrett-Byrne DA, Smith ND, Jamaluddin MFB, Kahl RGS, et al. Phospho-heavy-labeled-spiketide FAIMS stepped-CV DDA (pHASED) provides real-time phosphoproteomics data to aid in cancer drug selection. Clin Proteomics. 2022;19(1):48.

45. Murray HC, Miller K, Brzozowski JS, Kahl RGS, Smith ND, Humphrey SJ, et al. Synergistic targeting of DNA-PK and KIT signaling pathways in KIT mutant acute myeloid leukemia. Mol Cell Proteomics. 2023:100503.

46. Duchatel RJ, Mannan A, Woldu AS, Hawtrey T, Hindley PA, Douglas AM, et al. Preclinical and clinical evaluation of German-sourced ONC201 for the treatment of H3K27M-mutant diffuse intrinsic pontine glioma. Neurooncol Adv. 2021;3(1):vdab169.

47. Rose WC, and Wild R. Therapeutic synergy of oral taxane BMS-275183 and cetuximab versus human tumor xenografts. Clin Cancer Res. 2004;10(21):7413–7.

48. Tsherniak A, Vazquez F, Montgomery PG, Weir BA, Kryukov G, Cowley GS, et al. Defining a Cancer Dependency Map. Cell. 2017;170(3):564–76 e16.

49. Nunnery SE, and Mayer IA. Management of toxicity to isoform alpha-specific PI3K inhibitors. Ann Oncol. 2019;30 Suppl 10:x21–x6.

50. Nair AB, and Jacob S. A simple practice guide for dose conversion between animals and human. J Basic Clin Pharm. 2016;7(2):27–31.

51. Beekhof R, van Alphen C, Henneman AA, Knol JC, Pham TV, Rolfs F, et al. INKA, an integrative data analysis pipeline for phosphoproteomic inference of active kinases. Mol Syst Biol. 2019;15(5):e8981.

52. Weidner C, Fischer C, and Sauer S. PHOXTRACK-a tool for interpreting comprehensive datasets of post-translational modifications of proteins. Bioinformatics. 2014;30(23):3410–1.

53. Crul T, and Maleth J. Endoplasmic Reticulum-Plasma Membrane Contact Sites as an Organizing Principle for Compartmentalized Calcium and cAMP Signaling. Int J Mol Sci. 2021;22(9).

54. Steinberg SF. Structural basis of protein kinase C isoform function. Physiol Rev. 2008;88(4):1341–78.

55. Wager TT, Hou X, Verhoest PR, and Villalobos A. Central Nervous System Multiparameter Optimization Desirability: Application in Drug Discovery. ACS Chem Neurosci. 2016;7(6):767–75.

56. Kilburn LB, Kocak M, Decker RL, Wetmore C, Chintagumpala M, Su J, et al. A phase 1 and pharmacokinetic study of enzastaurin in pediatric patients with refractory primary central nervous system tumors: a pediatric brain tumor consortium study. Neuro Oncol. 2015;17(2):303–11.

57. DeWire M, Fuller C, Hummel TR, Chow LML, Salloum R, de Blank P, et al. A phase I/II study of ribociclib following radiation therapy in children with newly diagnosed diffuse intrinsic pontine glioma (DIPG). J Neurooncol. 2020;149(3):511–22.

58. Broniscer A, Baker JN, Tagen M, Onar-Thomas A, Gilbertson RJ, Davidoff AM, et al. Phase I study of vandetanib during and after radiotherapy in children with diffuse intrinsic pontine glioma. J Clin Oncol. 2010;28(31):4762–8.

59. Khan A, Gamble LD, Upton DH, Ung C, Yu DMT, Ehteda A, et al. Dual targeting of polyamine synthesis and uptake in diffuse intrinsic pontine gliomas. Nat Commun. 2021;12(1):971.

60. Persson ML, Douglas AM, Alvaro F, Faridi P, Larsen MR, Alonso MM, et al. The intrinsic and microenvironmental features of diffuse midline glioma: Implications for the development of effective immunotherapeutic treatment strategies. Neuro Oncol. 2022;24(9):1408–22.

61. Nagaraja S, Vitanza NA, Woo PJ, Taylor KR, Liu F, Zhang L, et al. Transcriptional Dependencies in Diffuse Intrinsic Pontine Glioma. Cancer Cell. 2017;31(5):635–52 e6.

62. Huang X, Liu G, Guo J, and Su Z. The PI3K/AKT pathway in obesity and type 2 diabetes. Int J Biol Sci. 2018;14(11):1483–96.

63. Hanker AB, Kaklamani V, and Arteaga CL. Challenges for the Clinical Development of PI3K Inhibitors: Strategies to Improve Their Impact in Solid Tumors. Cancer Discov. 2019;9(4):482–91.

64. Paugh BS, Broniscer A, Qu C, Miller CP, Zhang J, Tatevossian RG, et al. Genome-wide analyses identify recurrent amplifications of receptor tyrosine kinases and cell-cycle regulatory genes in diffuse intrinsic pontine glioma. J Clin Oncol. 2011;29(30):3999–4006.

65. Halvorson KG, Barton KL, Schroeder K, Misuraca KL, Hoeman C, Chung A, et al. A high-throughput in vitro drug screen in a genetically engineered mouse model of diffuse intrinsic pontine glioma identifies BMS-754807 as a promising therapeutic agent. PLoS One. 2015;10(3):e0118926.

66. Howell JJ, Hellberg K, Turner M, Talbott G, Kolar MJ, Ross DS, et al. Metformin Inhibits Hepatic mTORC1 Signaling via Dose-Dependent Mechanisms Involving AMPK and the TSC Complex. Cell Metab. 2017;25(2):463–71.

67. Labuzek K, Suchy D, Gabryel B, Bielecka A, Liber S, and Okopien B. Quantification of metformin by the HPLC method in brain regions, cerebrospinal fluid and plasma of rats treated with lipopolysaccharide. Pharmacol Rep. 2010;62(5):956–65.

68. Shen H, Yu M, Tsoli M, Chang C, Joshi S, Liu J, et al. Targeting reduced mitochondrial DNA quantity as a therapeutic approach in pediatric high-grade gliomas. Neuro Oncol. 2020;22(1):139–51.

69. Diabetes Prevention Program Research G. Long-term safety, tolerability, and weight loss associated with metformin in the Diabetes Prevention Program Outcomes Study. Diabetes Care. 2012;35(4):731–7.

70. Vinci M, Burford A, Molinari V, Kessler K, Popov S, Clarke M, et al. Functional diversity and cooperativity between subclonal populations of pediatric glioblastoma and diffuse intrinsic pontine glioma cells. Nat Med. 2018;24(8):1204–15.

71. Kline C, Jain P, Kilburn L, Bonner ER, Gupta N, Crawford JR, et al. Upfront Biology-Guided Therapy in Diffuse Intrinsic Pontine Glioma: Therapeutic, Molecular, and Biomarker Outcomes from PNOC003. Clin Cancer Res. 2022;28(18):3965-78.

72. Kawashima M, Hitomi Y, Aiba Y, Nishida N, Kojima K, Kawai Y, et al. Genome-wide association studies identify PRKCB as a novel genetic susceptibility locus for primary biliary cholangitis in the Japanese population. Hum Mol Genet. 2017;26(3):650–9.

73. Taylor KR, Barron T, Zhang H, Hui A, Hartmann G, Ni L, et al. Glioma synapses recruit mechanisms of adaptive plasticity. Preprint. 2021.

74. Kizub IV, Klymenko KI, and Soloviev AI. Protein kinase C in enhanced vascular tone in diabetes mellitus. Int J Cardiol. 2014;174(2):230–42.

75. Kalwa H, and Michel T. The MARCKS protein plays a critical role in phosphatidylinositol 4,5-bisphosphate metabolism and directed cell movement in vascular endothelial cells. J Biol Chem. 2011;286(3):2320–30.

