## Supplementary Figures for "PI3K/mTOR is a therapeutically targetable genetic dependency in diffuse intrinsic pontine glioma"

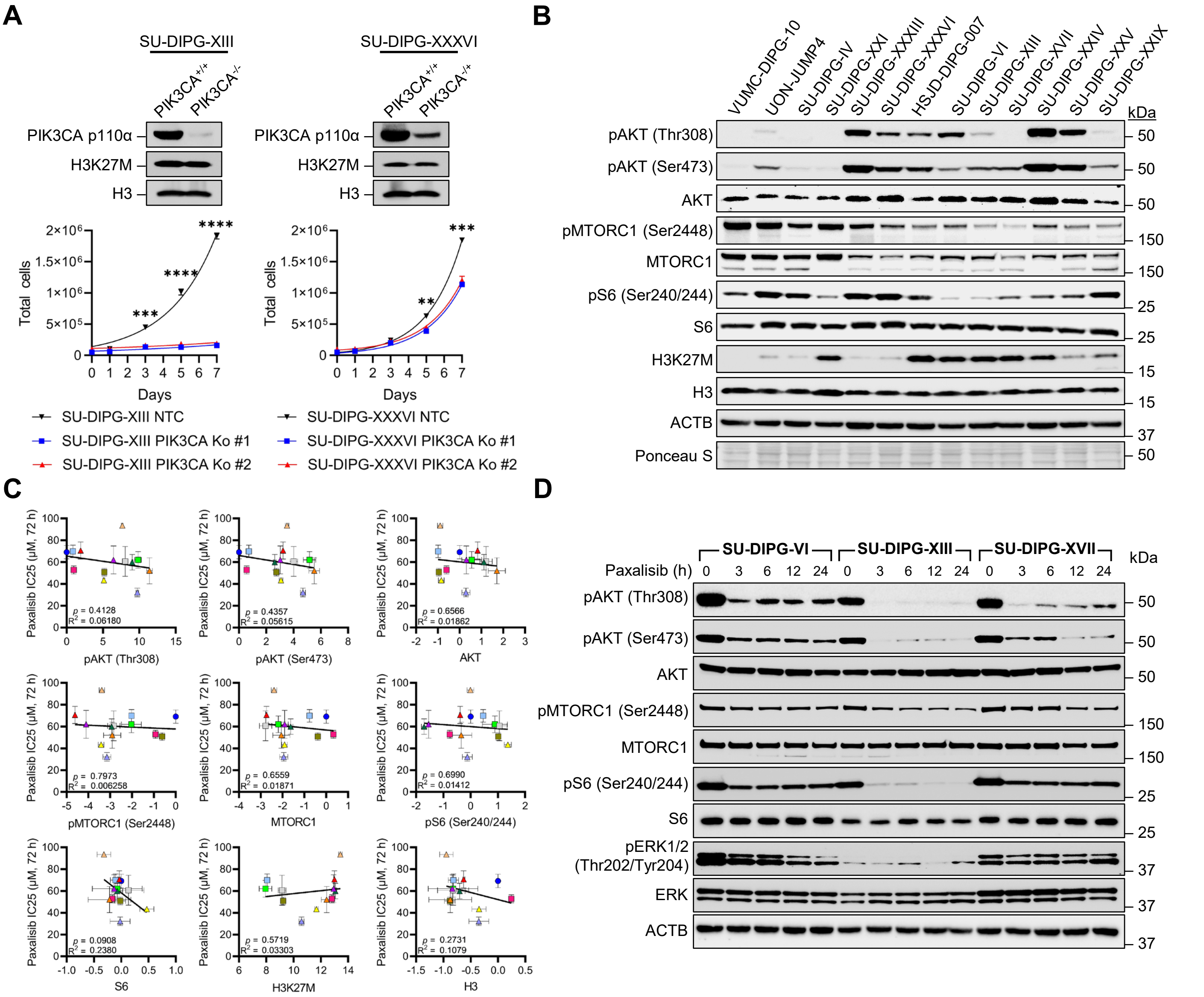


**Supplementary Figure S1: Expression and phosphorylation of PI3K/Akt/mTOR signaling proteins does not predict sensitivity to with paxalisib in vitro.** (A) Growth rates of SU-DIPG-XXXVI and SU-DIPG-XIII cells harboring *PIK3CA* knockout. (B) Protein expression of key PI3K/Akt/mTOR signaling proteins in DIPG cell lines measured by Western blot (n=3, representative Western blots presented). (C) Densitometry of Western blot values normalized to VUMC10 and correlated to IC25 values for paxalisib (significance determined using Pearson's linear regression). (D) Protein phosphorylation of key PI3K/Akt/mTOR signaling proteins in DIPG cell lines (SU-DIPG-VI, SU-DIPG-XIII, SU-DIPG-XVII) treated with paxalisib over time (1 µM, n=3, representative Western blots presented).

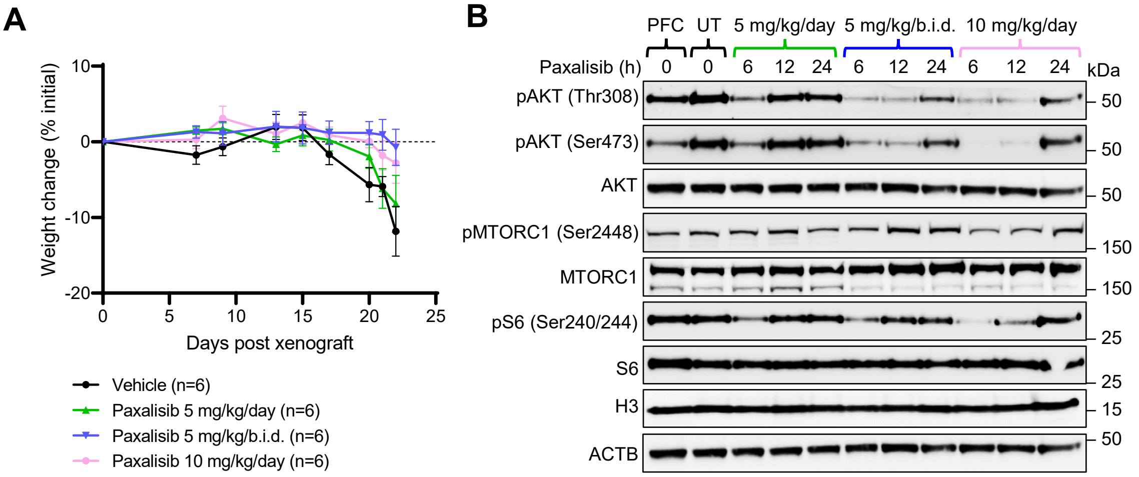


**Supplementary Figure S2: Paxalisib doe optimization maintains inhibition of PI3K/Akt/mTOR signaling.** (A) SU-DIPG-XIII-P* bearing xenograft mice weights were measure after treated with either vehicle, 5 mg/kg/day, 5 mg/kg/b.i.d., or 10 mg/kg/day treatment regime over time. (B) Tumors were resected from mice treated with vehicle, 6, 12 and 24 h, post treatment after 2 weeks of paxalisib treatment (vehicle, 5mg/kg/day, 5mg/kg/b.i.d., or 10mg/kg/day) and phosphorylation and expression of key PI3K/Akt/mTOR related signaling proteins measured by Western blot (n=3, representative Western blots presented).

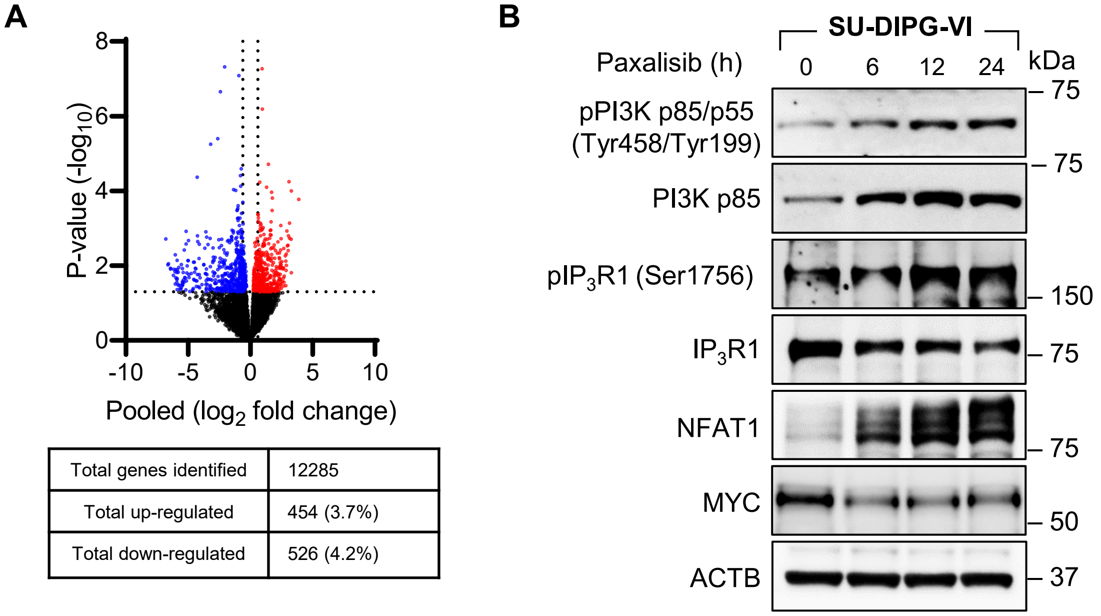


| | |
|---|---|
| Total genes identified | 12285 |
| Total up-regulated | 454 (3.7%) |
| Total down-regulated | 526 (4.2%) |

**Supplementary Figure S3: Analysis of gene expression following paxalisib treatment, validated via Western blotting.** (A) Bulk RNA Barcoding and sequencing (BRB-seq) of SU-DIPG-VI following paxalisib treatment (1 μM) for pooled data. (B) Validation of important oncogenes shown to be significantly modulated by paxalisib treatment at the protein level by Western blot (n=3, representative Western blots presented).

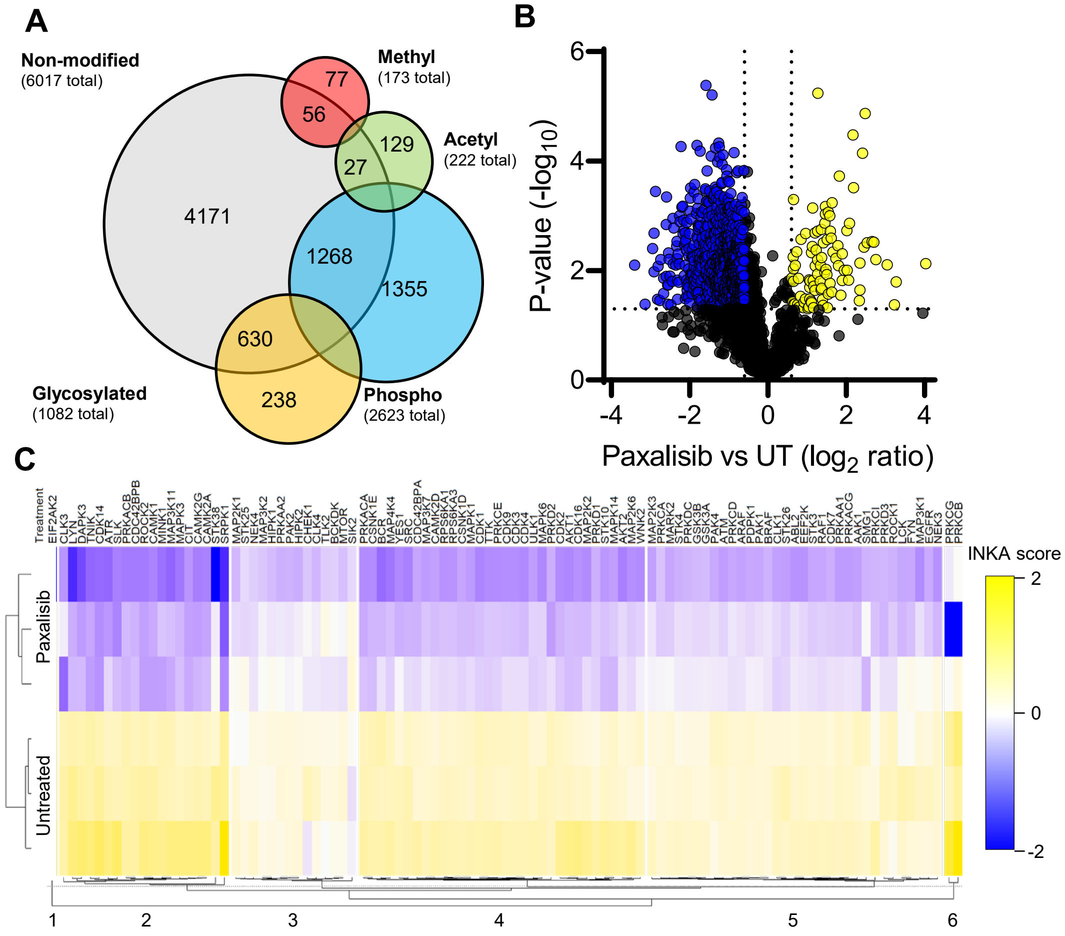


**Supplementary Figure S4: Phosphoproteomic profiling of paxalisib treated DIPG models.** (A) Phosphoproteomic profiling of SU-DIPG-XXXVI cells treated with 1 µM paxalisib for 6 h (n=3, biological triplicate). (B) Volcano plot of differentially phosphorylated proteins, (*p<0.05* log2 fold change <-0.6 and >0.6, n=3). (C) Integrated kinase expression analysis using Integrative inferred kinase activity (INKA) analysis of paxalisib treated cells (n=3).

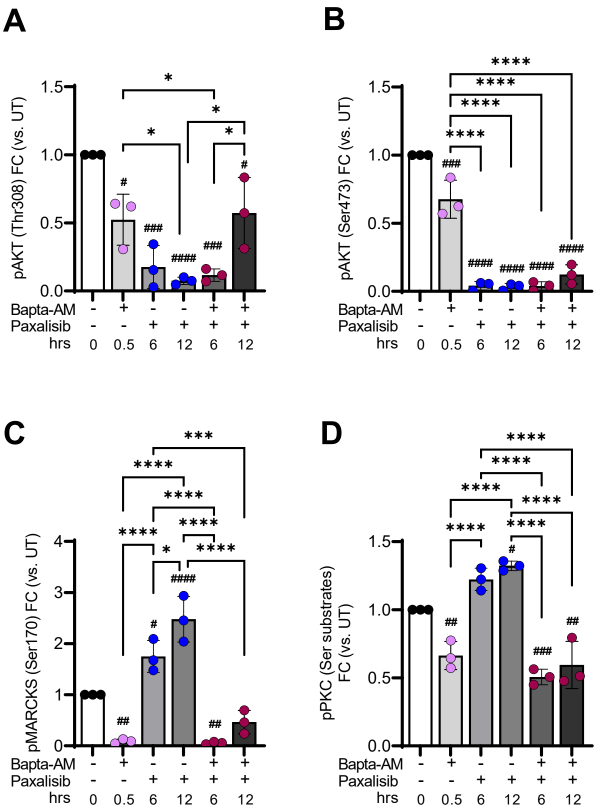


**Supplementary Figure S5: Increased calcium dependent PKC following paxalisib treatment.** Quantification of Western blot measurement of SU-DIPG-XIII cells treated with paxalisib or the calcium chelator BAPTA-AM (A) pAKT (Thr308), (B) pAKT(Ser473), (C) pMARCKS (Ser170) and (D) pSerPKC Substrates (n=3, one-way ANOVA, *$p<0.05$, **$p<0.01$, ***$p<0.001$, ****$p<0.0001$*).

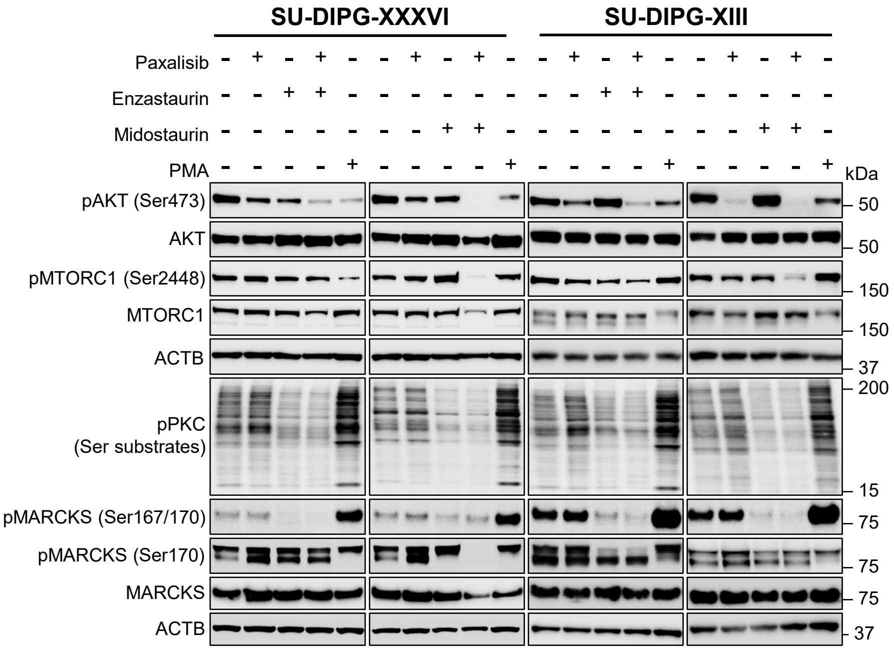


**Supplementary Figure S6: Assessment of PI3K- and PKC- related signaling following the combination of paxalisib with PKC inhibitors.** Protein expression and phosphorylation of key PI3K/Akt/mTOR and PKC signaling proteins was assessed after exposure to paxalisib (1 μM, 24 h), enzastaurin (5 μM, 24 h), midostaurin (5 μM, 24 h) and the PKC activator, PMA (1 μM, 24 h) (n=3, biological replicates, representative Western blots presented).

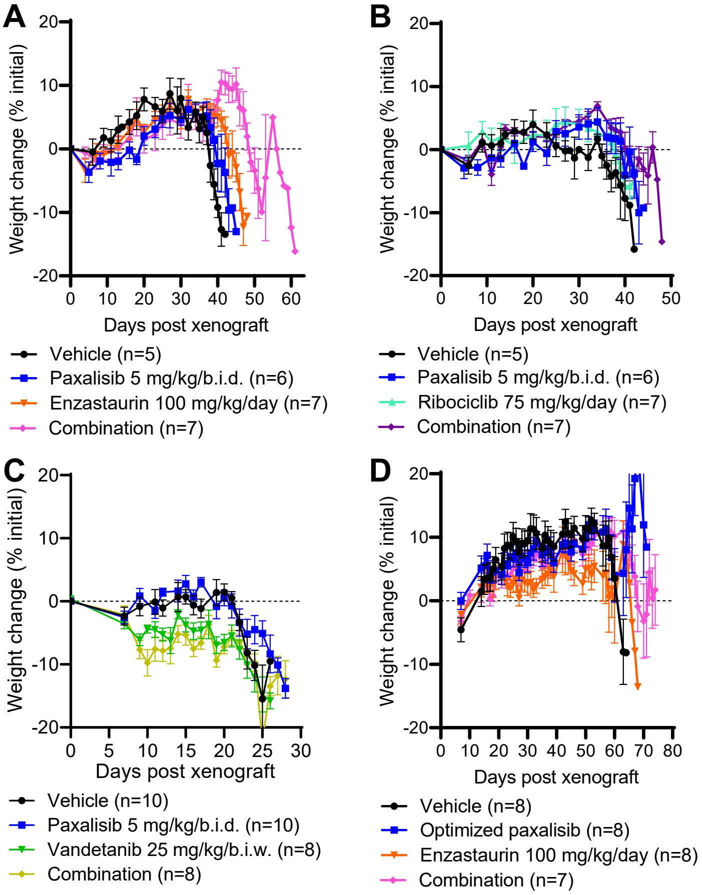


**Supplementary Figure S7: DIPG xenograft mouse weight following treatment with paxalisib alone and in combination with FDA approved therapies predicted to synergise by phosphoproteomic profiling.** Mouse weights recorded for SU-DIPG-XIII-P* xenograft models, following treatment with vehicle, paxalisib (5 mg/kg/b.i.d.) alone and combined with (A) enzastaurin (100 mg/kg/day), (B) ribociclib (75 mg/kg/day) or (C) vandetanib (25 mg/kg/b.i.w.). (D) RA-055 xenograft mouse model weights, following treatment with vehicle, optimized paxalisib (5 mg/kg/b.i.d., + 175 mg/kg/day metformin) in combination with enzastaurin (100 mg/kg/day).
